# A clickable photosystem I, ferredoxin, and ferredoxin NADP^+^ reductase fusion system for light-driven NADPH regeneration

**DOI:** 10.1101/2022.12.22.519867

**Authors:** Hitesh Medipally, Marvin Mann, Carsten Kötting, Willem J. H. van Berkel, Marc M. Nowaczyk

**Affiliations:** Plant Biochemistry, Faculty of Biology and Biotechnology, Ruhr University Bochum, Universitätsstr. 150, 44801 Bochum, Germany; Department of Biophysics, Faculty of Biology and Biotechnology, Ruhr University Bochum, Universitätsstr. 150, 44801 Bochum, Germany; Center for Protein Diagnostics (PRODI), Biospectroscopy, Ruhr University Bochum, Gesundheitscampus 4, 44801 Bochum, Germany; Laboratory of Food Chemistry, Wageningen University, Bornse Weilanden 9, 6708 WG Wageningen, The Netherlands; Department of Biochemistry, University of Rostock, Albert-Einstein-Str. 3, 18059 Rostock, Germany

**Author notes:** Corresponding author: Marc M. Nowaczyk.

**Keywords:** Photosystem I, Immunity protein 7, Ferredoxin, Ferredoxin NADP^+^ reductase, NADP^+^ photoreduction, NADPH, Fusion protein

## Abstract

Photosynthetic organisms like plants, algae, and cyanobacteria use light for the regeneration of dihydronicotinamide dinucleotide phosphate (NADPH). The process starts with the light-driven oxidation of water by photosystem II (PSII) and the released electrons are transferred via the cytochrome *b_6_f* complex towards photosystem I (PSI). This membrane protein complex is responsible for the light-driven reduction of the soluble electron mediator ferredoxin (Fd), which passes the electrons to ferredoxin NADP^+^ reductase (FNR). Finally, NADPH is regenerated by FNR at the end of the electron transfer chain. In this study, we established a clickable fusion system for in vitro NADPH regeneration with PSI-Fd and PSI-Fd-FNR, respectively. For this, we fused immunity protein 7 (Im7) to the C-terminus of the PSI-PsaE subunit in the cyanobacterium *Synechocystis* sp. PCC 6803. Furthermore, colicin DNase E7 (E7) fusion chimeras of Fd and FNR with varying linker domains were expressed in *E. coli*. Isolated Im7-PSI was coupled with the E7-Fd or E7-Fd-FNR fusion proteins through high-affinity binding of the E7/Im7 protein pair. The corresponding complexes were tested for NADPH regeneration capacity in comparison to the free protein systems demonstrating the general applicability of the strategy.

## Introduction

Photosystem I (PSI) is involved in light-driven electron transfer in photosynthetic organisms. It is regarded as one of the strongest biological reductants with high quantum efficiency and exceptional negative redox potential, which is sufficient to provide electrons to various intrinsic oxidoreductases involved in different metabolic processes of the cell, like carbon fixation or nitrogen assimilation.^[1–3]^ Due to the natural abundance and stability of PSI, development of biohybrid solar devices using PSI for efficient conversion of solar energy into chemical energy is a promising strategy.^[4–6]^ However, achieving direct electron transfer from PSI to a catalyst of interest is challenging due to its complex architecture and lack of cofactor exposure on the protein surface.^[7]^ These features distinguish PSI from other single-molecule photosensitizers, which can act as direct reductants for enzymes.^[8]^ The native and mobile redox mediator between PSI and many intrinsic oxidoreductases is ferredoxin (Fd).^[1]^ Therefore, electron transfer is a diffusion dependent process and the rate of electron supply depends on the interaction frequency of three components (PSI, Fd, and the terminal oxidoreductase). A tailored redox interface can facilitate the exploitation of the high electron yield of PSI to drive specific oxidoreductases of interest. Several strategies were proposed to achieve direct electron transfer from PSI.^[9–11]^ One of the most commonly used strategies is tethering the redox partner (Fd, oxidoreductase, or metal catalyst) onto the extrinsic subunits of PSI to achieve molecular confinement. This approach enables a rapid and direct electron transfer from PSI to the redox partner and also helps in channelling electrons to the specific oxidoreductase of interest. For example, by tethering a thiol linker-containing hydrogenase on a PSI-Cytc*_6_* fusion module, the electron transfer rate was two times higher compared to the natural system (105 vs. 45 e^-^ PSI^-1^ s^-1^).^[9]^Several types of other PSI-tethering strategies were proposed in recent years, such as reconstitution of Fd-PsaE-fusions to PSI complexes that were previously stripped and devoid of extrinsic stromal subunits ^[2]^, recombination of the fusion to PsaE deletion mutants ^[12]^, and site-directed assembly of Pt nanoparticles onto PSI.^[13]^ Molecular tethering of redox partners was also reported previously by direct genetic fusion of a hydrogenase to the extrinsic subunits of PSI (PsaE, PsaD, PsaC) ^[10,14,15]^ as well as a direct genetic fusion of Fd with a hydrogenase ^[16]^. All these models represent the combinations of bipartite fusions, i.e., either Fd or an oxidoreductase fused to PSI or Fd fused to the oxidoreductase. We have developed a multipurpose PSI fusion system, which allows tethering of various mediators and catalytic modules such as Fd or Fd-FNR, aiming for tailored and efficient electron transfer from PSI to its redox partners. This fusion is based on the high-affinity interaction of the E7-Im7 system. DNase E7 (E7) is a cytotoxic protein produced in *E. coli* and other *Enterobacteriaceae* strains as a stress response ^[17,18]^, which degrades DNA unspecifically. To avoid degradation of its own genomic DNA, the host cell produces the counter “Immunity protein 7” (Im7), which strongly binds to E7 and inhibits its activity. ^[19,20]^ These two proteins associate with high affinity (K_D_ = 10^-14^ to 10^-17^ M) and the protein couple can be used for highly efficient protein purification by affinity chromatography. ^[21]^ The cytotoxic activity of E7 can be suppressed by replacing a histidine at position 120 with alanine (H120A), allowing the safe recombinant expression of E7 in *E. coli* expression systems.^[22]^

In this study, Im7-modified PSI was generated and isolated from *Synechocystis* sp. PCC 6803 (hereafter *Synechocystis*) and combined with the recombinant chimeric protein constructs E7-Fd and E7-Fd-FNR as a proof of concept. Both model variants were tested based on light-induced dihydronicotinamide dinucleotide phosphate (NADPH) formation and compared with the free protein system.

## Results and Discussion

### Characterization of PSI-Im7-Histag

In this study, the N-terminus of Im7 was fused with the C-terminus of the PSI subunit PsaE. From a structural point of view, the PsaE subunit of PSI is exposed to the stroma and its C and N-terminus are freely accessible.^[7]^ For this reason, and the fact that the least physiological consequences are exhibited by PSI lacking the PsaE subunit ^[23]^, PsaE was chosen as the optimal subunit location for fusion with Im7. The fusion does not influence the growth rate of the PSI-Im7 mutant under provided growth conditions (Figure S1), indicating that this modification has no impact on the physiology of the strain. Blue Native (BN)-PAGE analysis revealed that the C-terminally His-tagged PSI-Im7 complex was successfully isolated using immobilized metal affinity chromatography (IMAC). In this way, we purified an equivalent of 3.2 mg chlorophyll (Chl) of PSI-Im7 from thylakoid membranes corresponding to the total Chl amount of 71 mg. The isolated PSI-Im7 was mostly present in the trimeric form (~1000 kDa; Figure1A). The observed bands above 1000 kDa on BN-PAGE may correspond to larger PSI aggregates. SDS-PAGE subunit analysis (Figure1B) of the PSI-Im7 complex was also performed, and individual subunits of PSI-Im7 were identified based on the previously determined masses of subunits. The following subunits of PSI-Im7 were identified: PsaA, PsaB, PsaC, PsaD, PsaE-Im7, PsaF. As expected, only IM7 modified PsaE was identified in PSI-Im7, indicating its successful integration into PSI. Identification of PsaC and PsaD further indicate that fusion of Im7 to PsaE did not influence the assembly of the other extrinsic subunits. Furthermore, spectral analysis also revealed no significant differences between the light-absorbing properties of PSI-Im7 and PSI-Wt (Figure S2A): both PSI complexes showed absorption maxima at 453 nm and 665 nm. [24]

**Figure 1.**
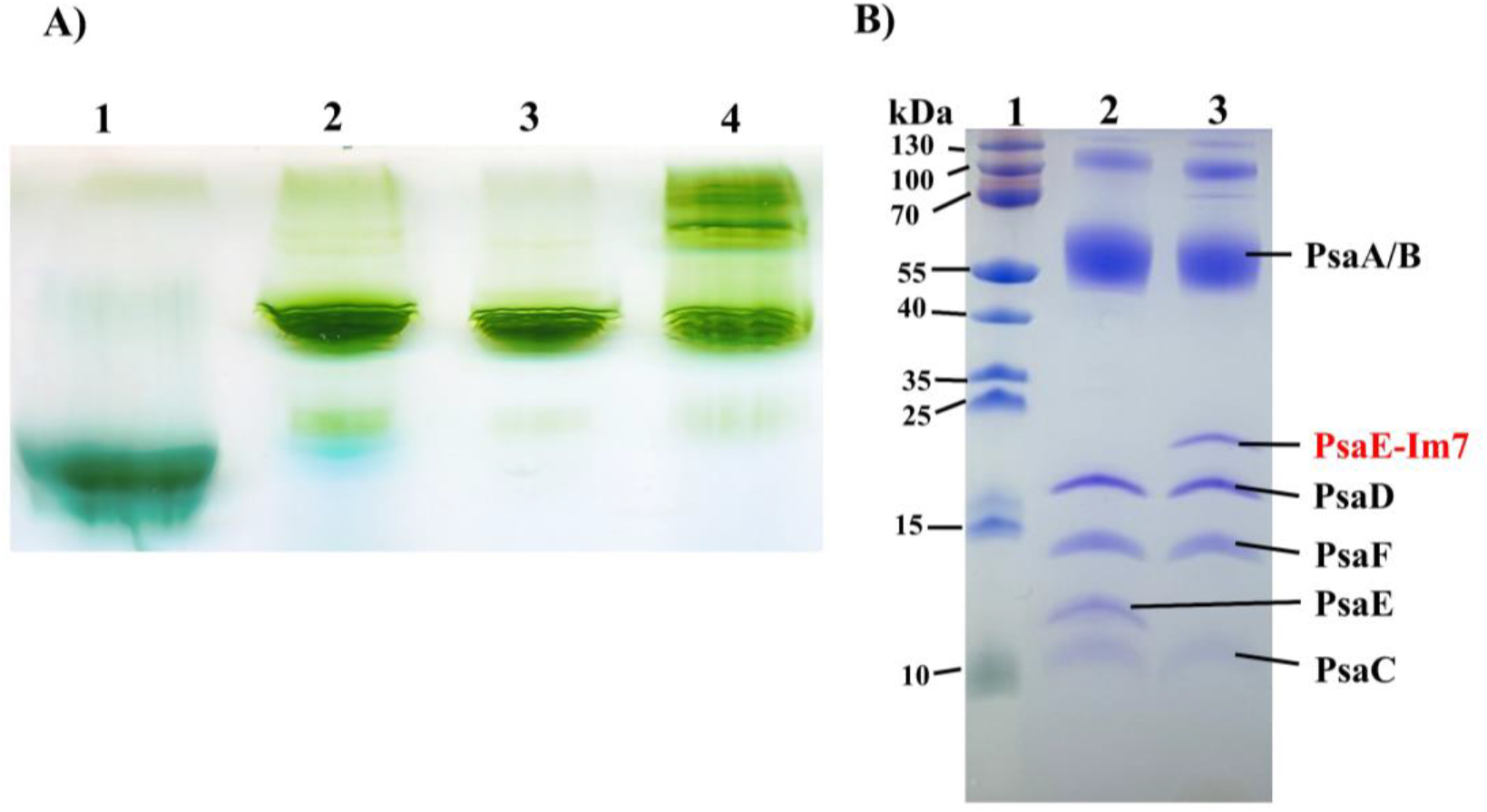
Biochemical characterization of the PSI-Im7-Histag complex. A) BN-PAGE-based analysis of the oligomeric state of PSI-Im7-Histag. 1: PSII monomer of *Thermosynechococcus vestitus* BP-1, 2: PSI-Wt trimer of *T. vestitus* BP-1, 3: PSI-Wt trimer of *Synechocystis*, 4: PSI-Im7-Histag complex of *Synechocystis* after IMAC purification. Equivalents of 3.5μg Chl were loaded onto each gel lane. B) SDS-PAGE analysis of PSI-Im7: 1: Protein marker (PageRuler Thermoscientific^™^). 2: PSI-Wt of *Synechocystis*, 3: PSI-Im7-Histag of *Synechocystis*. Equivalents of 2.5 μg Chl were loaded onto each gel lane.

The photoreduction rate of oxygen by isolated PSI-Im7 was comparable to that of PSI-Wt being1827 ± 21 μmol O_2_ mg Chl^-1^ mL^-1^ h^-1^ for PSI-Wt and 2038 ± 24 μmol O_2_ mg Chl^-1^mL^-1^ h^-1^ for PSI-Im7, respectively (Figure S2B). Thus, the PsaE-Im7 fusion does not influence PSI activity. The ability of PSI-Im7 to bind to DNase E7 and the photochemical activity of the complex was additionally verified by infrared difference spectroscopy. For this purpose, the Im7 tagged PSI was circulated over a chemically functionalized and E7-covered germanium internal reflection element (IRE).

Figure S3 shows the binding of PSI-Im7 on this surface and the stability of the complex during the subsequent washing step. In the next step, the surface with the immobilized PSI-Im7 complex was used to investigate the light-induced conformational change of the protein. For comparison, the light-induced difference spectrum was also measured in a transmission measurement for PSI in solution. In both setups a similar difference spectrum was obtained, indicating native complex formation and immobilization of PSI (Figure S4). Due to the thickness of the transmission cuvette, total absorption by water occurred in the range between 1680 cm^-1^ and 1600 cm^-1^ in the transmission measurement, while the attenuated total reflection (ATR) measurement provided an evaluable difference spectrum in this spectral range. Both spectra agree with measurements of PSI-Wt in the literature.^[25,26]^ According to Hastings et al. 2015, the most prominent bands can be assigned as follows: The negative band at 1700 cm^-1^ as well as the positive band at 1687 cm^-1^ originate from the keto C=O vibration of the A0 state of chlorophyll a. The positive band at 1755 cm^-1^ and the negative band at 1749 cm^-1^ can be assigned to the ester C=O vibration of the A0 state of chlorophyll a. The negative band at 1636 cm^-1^ corresponds to a C=C vibration of neutral phylloquinone (PhQ). The positive band at 1687 cm^-1^ and the negative band at 1670 cm^-1^ result from amide-I modes. In conclusion, the biochemical and spectroscopic analysis of PSI-Im7 did not reveal any major difference compared to PSI-Wt.

### Isolation and characterization of chimeric E7-Fd and E7-Fd-FNR fusion proteins

C-terminally His-tagged E7-Fd (EF) and E7-Fd-FNR (EFF) fusion proteins were produced in *E. coli* B121 Δ*iscR* and purified by IMAC. The chimeric fusion proteins consist of a flexible amino acid (AA) linker of GGSG motifs between individual domains. The purity of each chimeric fusion protein was checked by SDS-PAGE (Figures S5 and S6) and successful cofactor assembly was demonstrated by UV-Vis absorption spectroscopy (Figure 2). The EF chimeras (linker length variations of 5, 10, and 15 AA) showed typical Fd absorption properties with spectral maxima at 330 and 420 nm (Fig. 2A and 2C). The incorporation of the Fe-S cofactor was investigated by determining the ratio of 420/278 nm, which amounted to 0.59 for Fd and ~0.44 for EF (mean value of all variants), respectively. The former value was similar to the expected ratio of fully assembled Fd (0.56) reported previously ^[2]^, while the reduced ratio observed for the EF fusion proteins can be explained by the presence of the additional E7 domain, which increases the absorption at 280 nm but not at 420 nm. For example, the calculated extinction coefficient of EF10 at 280 nm is 24.32 mM^-1^ cm^-1^ and that of the E7 domain alone is 12.49 mM^-1^ cm^-1^. As a result the normalized absorption spectra of both EF and EFF chimeras (linker length variations between Fd and FNR was 5, 10, 15 AA) are lower than those of the free Fd and FNR proteins (Figure 2C and 2D). Additionally, minor protein impurties may have contributed to the absorbance values of the EF and EFF chimeras at 280 nm. The absorption spectrum of FNR (Figure 2D) represented the typical pattern of a flavoprotein with absorption maxima at 394 nm and 459 nm, as well as a shoulder at 484 nm^[27,28]^ and the typical absorption maxima of Fd were observed at 420 and 330 nm. In case of EFF chimeras (linker length variations between Fd and FNR of 5, 10, and 15 AA), the absorption maxima at 394 nm and 459 nm were observed (Figure 2D), indicating the assembly of the flavin adenine dinucleotide cofactor into the FNR domain of EFF. The 459/280 nm ratios for FNR and EFF were 0.29 and ~0.19 (mean value of all variants), respectively, which can be explained by the presence of the E7 and Fd domains in the fusion protein. Due to strong absorption of the EFF chimeras at 394 and 459 nm, the absorption peak at 420 nm related to the Fe-S cluster of the Fd domain was masked. However at 330 nm, the absorption of the EFF chimeras was higher compared to FNR, consistent with the assembly of the Fe-S cluster into the Fd domains of EFF chimeras.

**Figure 2.**
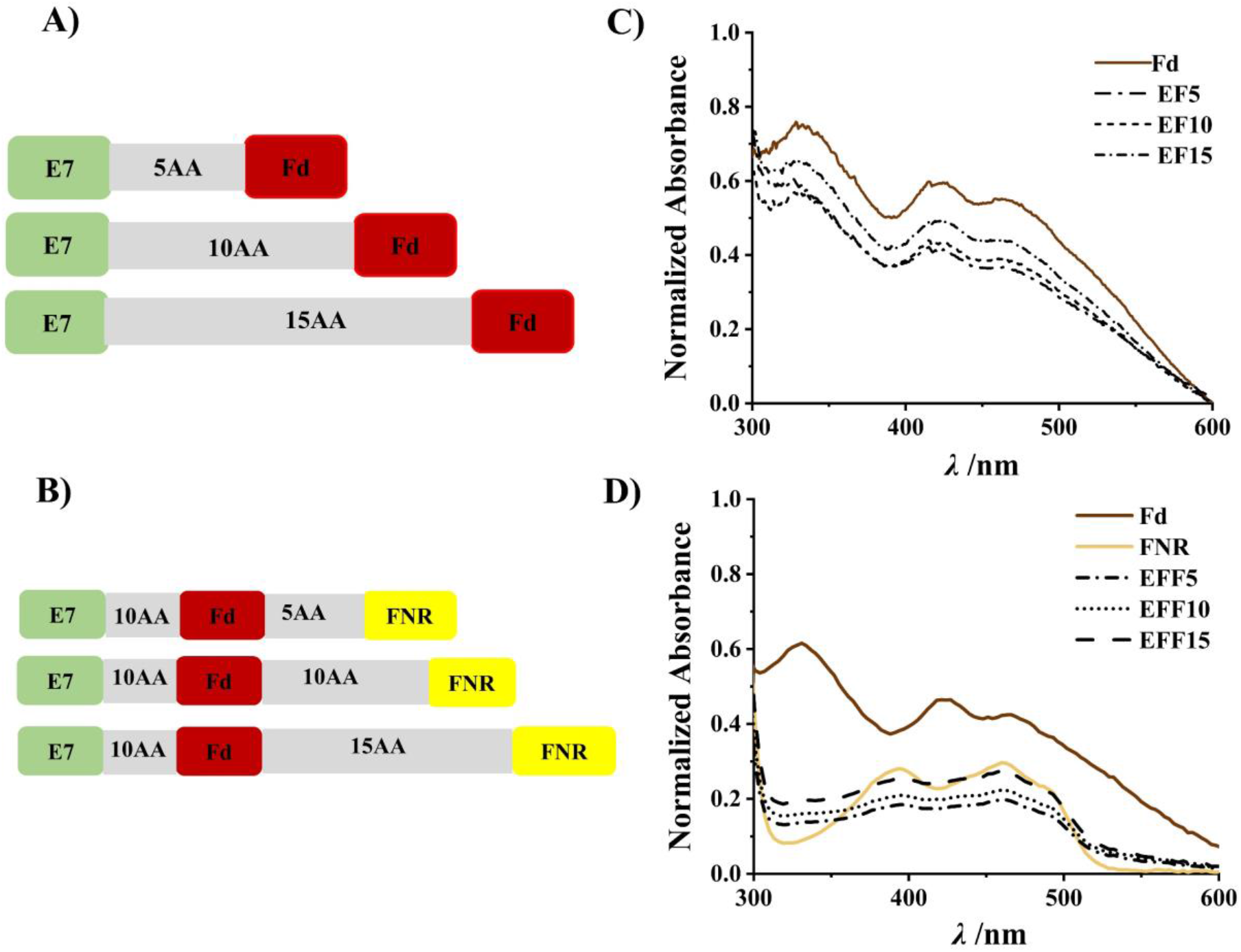
Design and spectral characterization of E7-Fd (EF) and E7-Fd-FNR (EFF) chimeras. A) Genetic design of E7-Fd chimeras with respective linker lengths of 5, 10, and 15 AA. B) Genetic design of E7-Fd-FNR with a constant linker length of 10 AA between E7 and Fd and varying linker lengths of 5, 10, and 15 AA between Fd and FNR. C) UV-Vis absorption spectra of His-tag-purified Fd and EF chimeras with absorption normalized to 278 nm and absorption maxima at 420 and 330 nm. D) UV-Vis absorption spectra Fd (brown), FNR (yellow) and EFF chimeras (black) with absorption normalized to 278 nm and absorption maxima at 394, 459 nm. Experiments were conducted in triplicates and the data represents mean values.

### Coupling of EF and EFF chimeras to PSI-Im7

The strong affinity of E7 for Im7 should promote coupling of EF and EFF variants to PSI-Im7 even at very low concentrations, avoiding heterogeneity of the in vitro system. This feature will also provide an advantage in the case of in vivo implementation of the PSI fusions or in the establishment of other in vivo fusion systems. Nonetheless, to ensure that every protomer in trimeric PSI-Im7 binds chimeric EF/EFF variants, we used a 1:8 molar ratio of PSI-Im7 (per protomer) and EF/EFF. The coupled complexes and excess of EF/EFF were separated by sucrose density gradient centrifugation (SDGC). The large differences in molecular weight between the coupled complexes (~1100 kDa) and EF/EFF (<100 kDa) chimeras allow easy separation via SDGC (Figure S7). The homogeneity and integrity of PSI chimeras after SDGC separation was confirmed by SDS-PAGE (Figure 3) and BN-PAGE analysis (Figures S8, S9).

**Figure 3.**
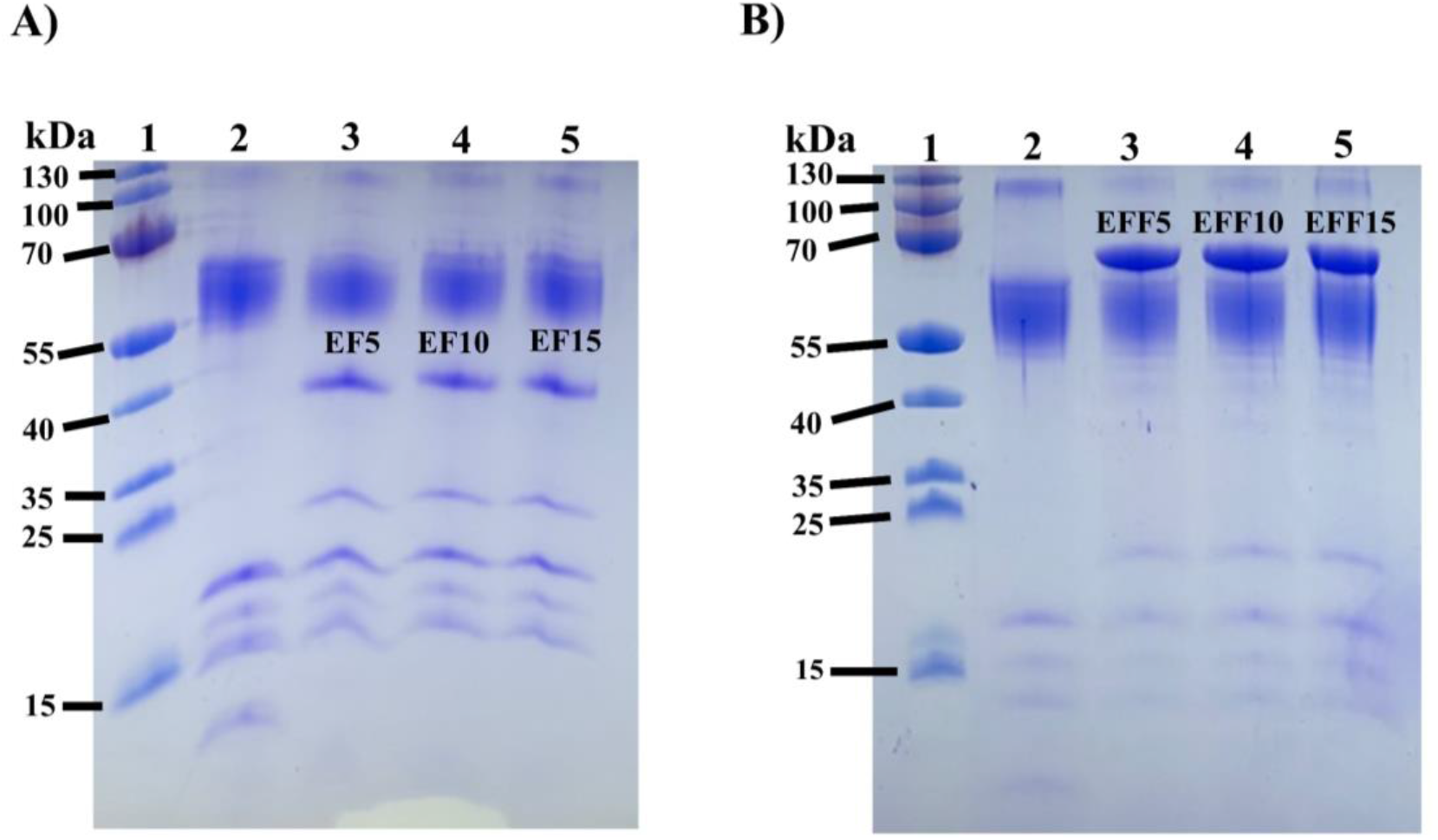
SDS-PAGE analysis of PSI-Fd (PSI-EF) and PSI-Fd-FNR (PSI-EFF) chimeras after SDGC separation. A) SDS-PAGE analysis of PSI-EF chimeras. 1: Protein marker (PageRuler, Thermoscientific^™^) with masses of individual protein bands indicated on the left; 2: PSI-Wt; 3: PSI-EF5; 4: PSI-EF10; 5: PSI-EF15. B) SDS-PAGE analysis of PSI-EFF chimeras. 1: Protein marker (PageRuler, Thermoscientific^™^) with masses of individual protein bands indicated on the left; 2: PSI-Wt; 3: PSI-EFF5; 4: PSI-EFF10; 5: PSI-EFF15. An equivalent of 2.5 μg Chl was loaded onto each gel lane.

### Light-induced NADP^+^ reduction of PSI-Fd chimeras

In vitro light-induced electron transfer of PSI-Fd (PSI-EF) chimeras with variable linkers (PSI-EF5, PSI-EF10, PSI-EF15) was characterized by artificial reconstitution of the photosynthetic electron transport chain, including free FNR. Differences in the redox potentials between individual protein components allowed the successive electron transfer in a reconstituted cascade, i.e. from ascorbate to NADP^+^. In this assay, light-induced electron transfer rates of PSI-EF5, PSI-EF10, PSI-EF15 chimeras, and the free protein system (FPS1) were determined to be 2.05 ± 0.22, 2.49 ± 0.08, 1.51 ± 0.20 and 2.02 ± 0.29 μM NADPH min^-1^ (Figure 4). With PSI-EF10, the rate of light-induced NADP^+^ reduction compared most favorably with that of FPS1, strengthening the assumption that the optimal linker length between E7-Fd would be around 10 AA. Moal and coworkers reported that the NADP^+^ reduction rate of a PSI-PsaE-Fd coupled construct with a linker length of 10 AA between PSI-PsaE and Fd outperformed the FPS by 1.12-fold.^[12]^ To avoid too strong association between PSI and Fd, which resulted in absence of catalytic activity, they substituted Arg39 of PsaE with Gln39. ^[12]^ In the case of PsaE-Im7, the strong association of Fd with PSI was not observed. This might be due to an extra distance or structural change created by the presence of E7-Im7. With a short linker (PSI-EF5), the initial rate of light-induced NADP^+^ reduction was about equal to the rate obtained for the free protein system (Figure 4). However, in the case of PSI-EF15, the light-induced NADP^+^ reduction was significantly less efficient than with FPS1 (Figure 4), suggesting that the long linker and additional space created by E7-Im7 made this fusion system too flexible. The results obtained are in agreement with those of Yacoby et.al, who constructed a PSI-based electron transport chain with Fd-hydrogenase fusions for the anaerobic production of hydrogen gas. ^[29]^ The highest electron transfer rate of these Fd-hydrogenase fusions was achieved with a linker length of 25 AA. However, with 30 AA, the hydrogen production dropped significantly, which could be explained by diffusion-based constraints in the interaction of Fd with PSI and hydrogenase.

**Figure 4:**
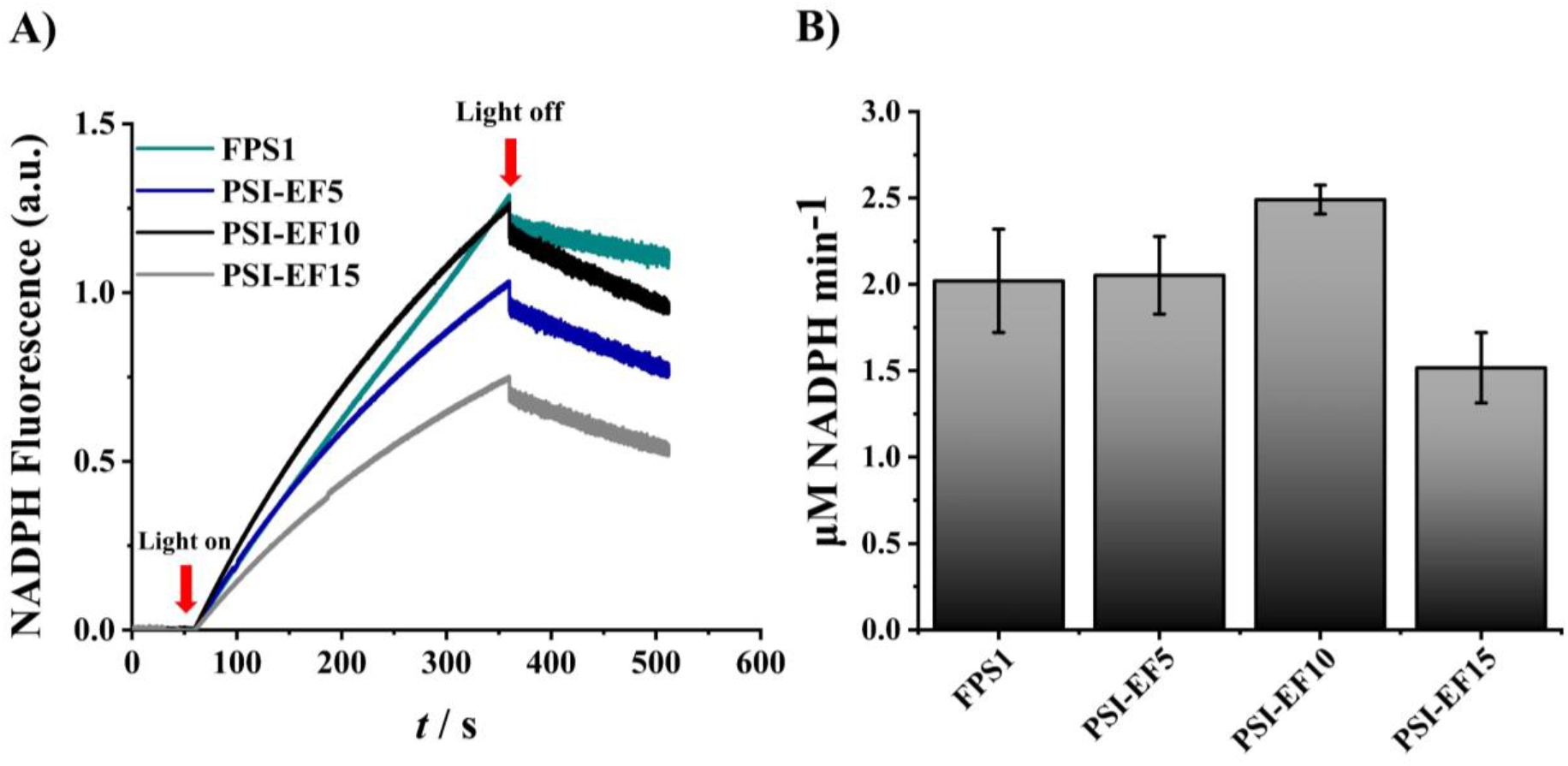
Light-induced NADP^H^ production by PSI-Fd chimeras. A) Light-induced NADPH formation of FPS1 (cyan), PSI-EF5 (blue), PSI-EF10 (black), and PSI-EF15 (gray) was measured by NADPH fluorescence. The concentration of each fusion protein was 0.25 μM and the concentration of free FNR was 0.75 μM. FPS1 contained 0.25 μM PSI, 0.25 μM Fd, and 0.75 μM FNR. The total measuring time was 8 min with a dark (1 min)-light (5 min)-dark (2 min) cycle. All the experiments were performed in 1.5 mL reaction buffer (50 mM Tricine pH 8, 30 mM NaCl, 4 mM NaAsc, 100 μM DCPIP, 2.5 μM Cytc*6*, 5 mM MgCl_2_ (added only in the case of FPS1),1 mM NADP^+^, 0.03% DDM) by artificial reconstitution of the photosynthetic electron transport chain from ascorbate to NADP^+^. B) Initial rates of light-induced NADPH formation rates. Experiments were conducted in triplicates, and the data points represent the mean values with standard deviation.

### Light-induced NADP^+^ reduction of PSI-Fd-FNR chimeras

The PSI-Fd-FNR (PSI-EFF) chimeras were also characterized by artificial reconstitution of the photosynthetic electron transport chain from ascorbate to NADP^+^ and the NADP^+^ photoreduction rates were compared with a free protein system (FPS2). The PSI-EFF linker variants (PSI-EFF5, PSI-EFF10 and PSI-EFF15) showed different electron transfer rates, as illustrated in Figure 5. The initial rates of NADP^+^ photoreduction of FPS2, PSI-EFF5, PSI-EFF10 and PSI-EFF15 were 5.30 ± 0.62, 3.60 ± 0.21, 1.63 ± 0.15, 1.07 ± 0.03 μM NADPH min^-1^, respectively. In FPS2, a typical electron supply from PSI via Fd to FNR is expected, i.e. the simultaneous interchange of Fd_ox_ with Fd_red_ at the Fd-binding site of FNR for reduction of NADP^+^ by FNR.^[30]^ In case of the fusion constructs, two-sided tethering of Fd may have introduced steric hindrance in the electron transfer. Nevertheless, PSI-EFF5 activity was only slightly less compared to the activity of FPS2. With longer linker lengths, a decrease in NADP^+^ photoreduction rate was observed. We speculate that the wide distance introduced by the linker may have created a lag in the interaction of Fd with FNR. As a result, PSI-EFF5 showed a higher NADP^+^ photoreduction rate than PSI-EFF10 and PSI-EFF15, respectively. The functional behavior of each fusion complex might be explained more evidently in future work by building structural models and applying molecular dynamics simulations ^[31]^. Furthermore, the NADPH formation rates of the tripartite system was observed to be faster compared to the bipartite system. The relatively faster NADPH formation rate in the tripartite system could be explained by the faster electron transfer rates between PSI, Fd and FNR in the initial phase of the reaction (Figure 5). However, in the mid and final phase of the reaction, the higher NADPH fluorescence rate can be attributed to accumulated NADPH and its slow consumption by FNR. This is due to lower FNR concentration in the tripartite system sample (0.25 μM) compared to the bipartite system sample (0.75 μM), which is further confirmed by fast consumption of NADPH in the dark in case of the bipartite system (Figure 4 and 5).

**Figure 5:**
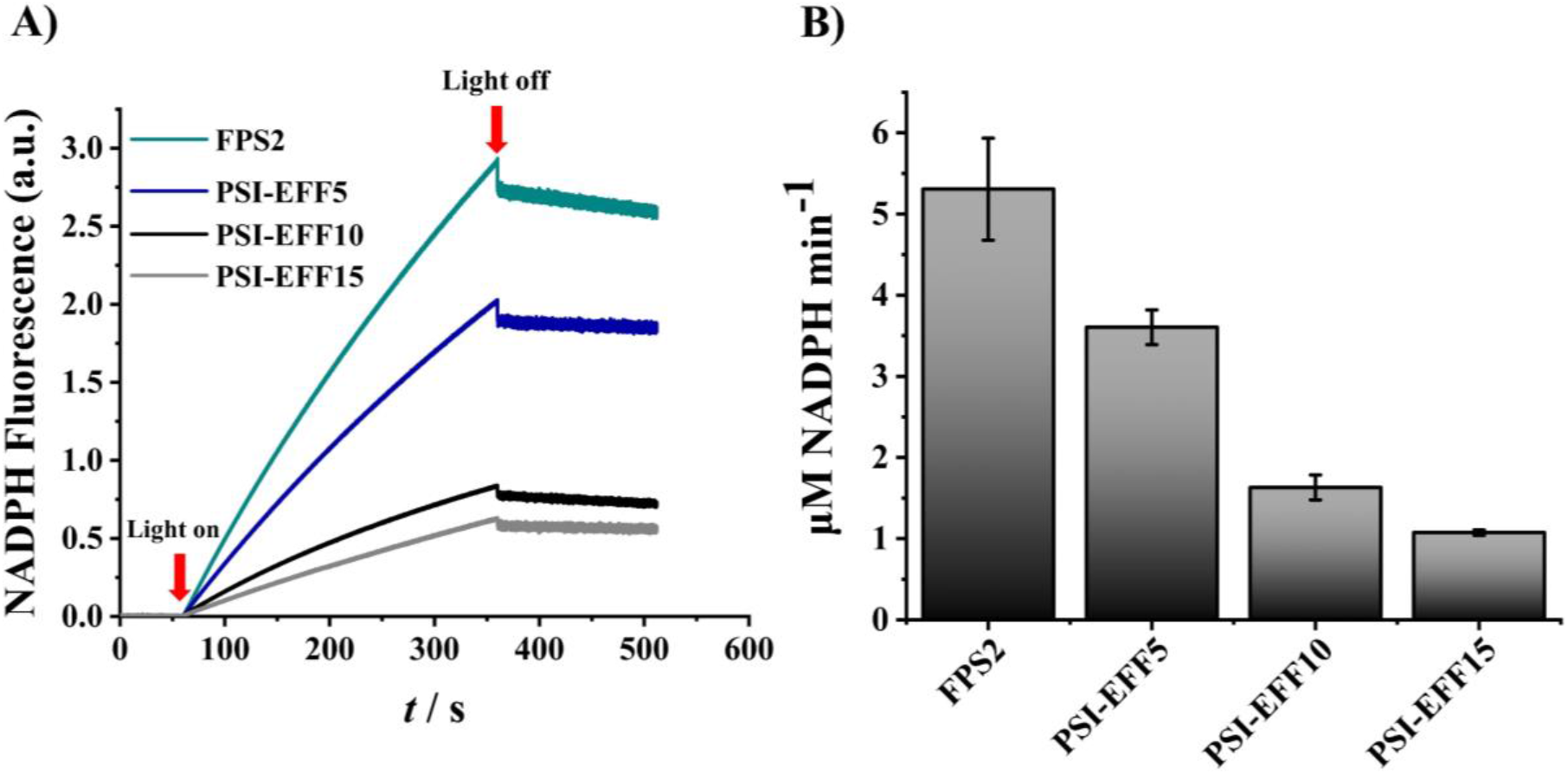
Light-induced NADPH production by PSI-Fd-FNR chimeras. A) Light-induced NADPH formation of FPS2 (cyan), PSI-EFF5 (blue), PSI-EFF10 (black), and PSI-EFF15 (gray) was measured by NADPH fluorescence. The concentration of each fusion protein was 0.25 μM. FPS2 contained 0.25 μM PSI, 0.25 μM Fd, and 0.25 μM FNR. The total measuring time was 8 min with a dark (1 min)-light (5 min)-dark (2 min) cycle. All the experiments were performed in 1.5 mL reaction buffer (50 mM Tricine pH 8, 30 mM NaCl, 4 mM NaAsc, 100 μM DCPIP, 2.5 μM Cytc*6*, 5 mM MgCl_2_ (added only in the case of FPS1),1 mM NADP^+^, 0.03% DDM) by artificial reconstitution of photosynthetic electron transport chain from ascorbate to NADP^+^. B) Initial rates of light-induced NADPH formation. Experiments were conducted in triplicates, and the data represent the mean value with standard deviation.

### Light-induced electron transfer in various modified PSI fusion complexes

Several PSI fusion models proposed are tabulated below (Table 1). Although each PSI fusion model was developed in combination with different enzymes or mediators and assessed at different reaction conditions, they still can be compared based on electron transfer efficiency, i.e. number of electrons released out of light-excited PSI per second (e PSI^-1^ s^-1^). For this, the PSI amount was calculated by considering each PSI monomer consisting of 95 chlorophyll molecules ^[32]^. Among the various PSI fusion models, PSI was modified, particularly to drive hydrogen production in vitro, by using platinum or a hydrogen producing enzyme (FeFe or NiFe type hydrogenases) as a catalytic module. Greenbaum and coworkers pioneered the photocatalytic production of hydrogen by developing PSI-metal nanoparticles with direct precipitation of charged Pt ligands and colloidal Pt onto isolated PSI or chloroplasts, respectively ^[33,34]^. These models are catalytically less active (Table 1), and deposition of Pt in proximity to the PSI active site was not regulated. Another approach was the development of a molecular wire, which connects the terminal F_B_ cluster of PSI with Pt or hydrogenase for direct electron transfer via a hexane-1,6-dithiol linker integrated into PSI by ligand rescue. In this procedure, recombinant PsaC in which the cysteine residue at position 13 was replaced by glycine (C13G) was prepared from *E. coli* and reconstituted artificially with Fe-S clusters. This enabled the tethering of one end of the thiol linker to the terminal F_B_ cluster and the other to the catalyst. Further, this artificially reconstituted PsaC subunit with the catalyst attached was later integrated into PSI ^[9,35,36]^. With this approach, the catalyst was kept near the active site of PSI and the electron transfer rates obtained with this system were among the highest reported in the literature. However, this method, which employs synthetic thiol linkers, is not compatible with in vivo approaches because it cannot be applied to whole cells. An alternative approach is based on the in vitro reconstitution of a chimeric hydrogenase with PSI, which was isolated from a *ΔPsaE* mutant.^[15]^ In this approach, amino acid-based linkers were used to fuse the hydrogenase with PsaE, which enables in principle also in vivo applications. Indeed, it was shown that the photoautotrophic growth of the PSI-*ΔPsaE* mutant was not affected.^[23]^ However, the direct electron transfer towards HydA1 coupled to PSI via PsaE was observed to have very low electron transfer rates (Table 1). Recently, Appel et al. and Kanygin et al. showed successful hydrogen production by in vivo PSI modification via direct electron transfer by genetic fusion of the hydrogenase to one of the extrinsic subunits (PsaC, PsaD) of PSI.^[10,14]^ However, in both models, physiological constraints were noticed. For instance, in PSI-PsaD-Hox-YH-mutant from the cyanobacterium *Synechocystis*, binding of the PsaD-fusion displaces the PsaC subunit of PSI, and it is assumed that electrons are accepted from the F_x_ cluster.^[14]^ Moreover, the PSI-PsaC-HydA fusion from the eukaryotic microalgae *Chlamydomonas reinhardtii* showed reduced Fd binding ability with PSI up to 50%.^[10]^ Therefore, modification of PsaC or PsaD with hydrogenase is considered a strong mutation leading to poor accumulation of hydrogenase-modified PSI in cells.^[10]^ In case of PSI-Im7, the subunit composition and photoreduction activity of PSI are not affected. Also, within PSI-EFF chimeras, a decrease in linker length between Fd and FNR, proportionally increased the NADP^+^ photoreduction rate of PSI-EFF chimeras. The electron transfer within PSI-EFF can be further improved by redesigning the linker between Fd and FNR and predicting and testing the suitable catalyst orientation and interaction frequency between PSI, Fd and catalyst might improve the photoreduction rates of the fusion complex.

**Table 1:**
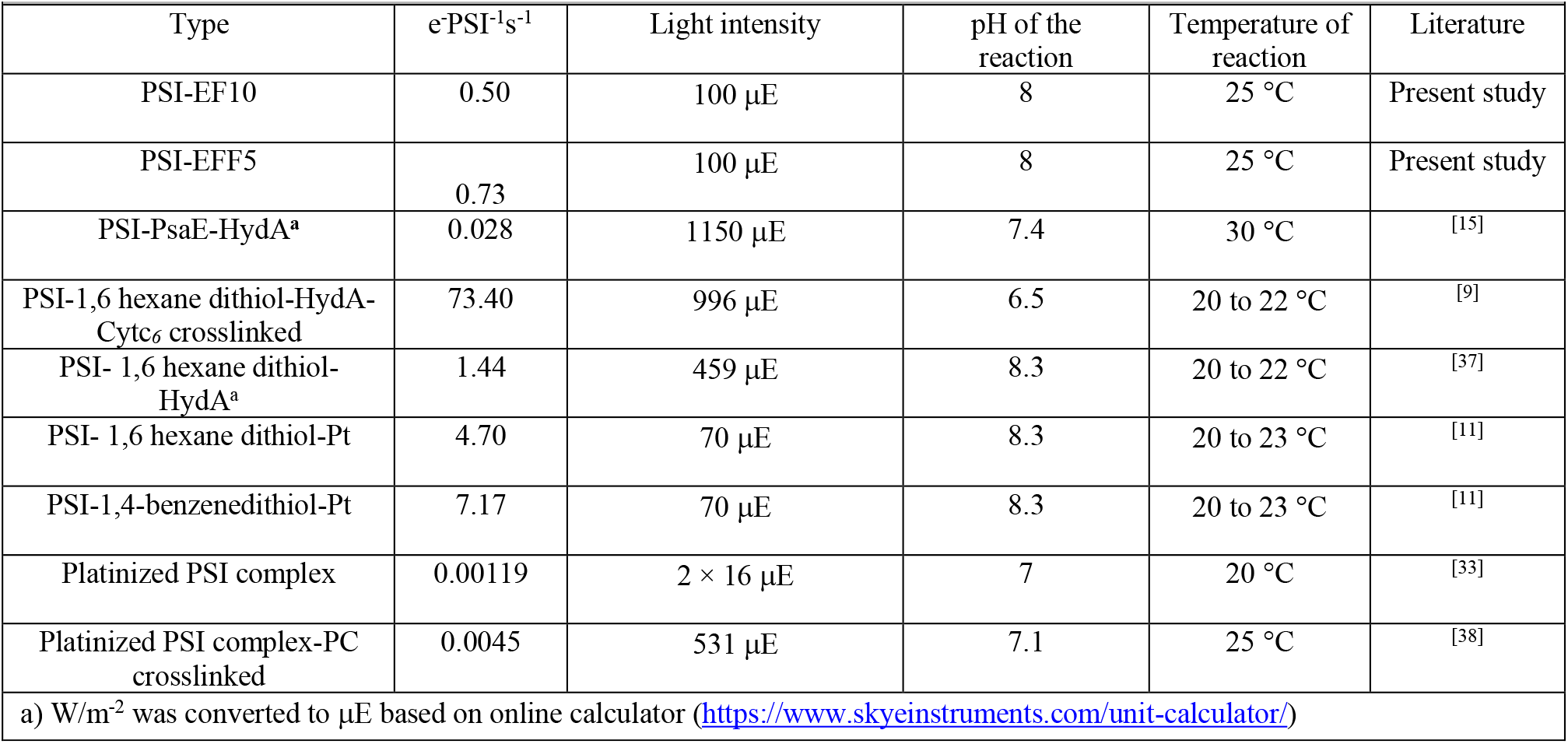
Light-induced electron transfer rate in modified PSI complexes

### Application of PSI-Im7 fusion model in development of next-generation photovoltaic cells

PSI has been extensively used as a photochemical module in the development of semi-artificial photosynthetic devices. ^[39–46]^ However, insulation of acceptor and donor side of PSI on the electrode surface to avoid charge recombination is challenging and was not realized in many of the PSI-electrode integration models. Due to the lack of insulation, reduced electron acceptors such as methyl viologen (MV) short-circuit the PSI electron input and output side. As a result, the photocurrent output of these models was compromised. In nature, PSI is incorporated into the thylakoid membrane in a defined orientation, preventing short-circuiting between the acceptor and donor side of PSI. In analogy with thylakoids, PSI was fabricated onto the electrode surface in an oriented manner via Langmuir Blodgett technology. ^[41]^ However, the dispersed and spatial distribution of PSI trimers achieved with Langmuir Blodgett technology on the electrode surface still contributes to short-circuiting effects. Recently, a significant improvement was made to reduce this by closing the gaps between the PSI trimers with PSI monomers to nullify the effect of spatial distribution. ^[47]^ On the other hand, to realize the complete potential of the electron transfer efficiency of PSI, a robust redox interface on both the donor and acceptor sides of PSI is essential. With lack of a proper electron acceptor, electrons from a photoexcited PSI F_B_ cluster back transfer to internal cofactors F_A_, F_X_, and A_1A_, successively ^[48]^, resulting in charge recombination. Alternatively, the electrons are delivered to an immediately available electron acceptor. In photosynthetic cells, oxygen is such an example. Its reduction results in the formation of reactive oxygen species (ROS) that cause photodamage. Development of a semi-artificial photosynthetic device, which addresses the effect of charge recombination within PSI and short-circuiting within the system can be potentially created by integration of PSI-EFF fusion complexes on the electrode via Langmuir Blodgett technology. Due to the built-in natural electron mediator within the PSI-EFF construct, the short-circuiting effect should be reduced, and the proximal presence of Fd near the PSI active site may avoid charge recombination within PSI. Another potential application of the PSI-EFF fusion model would be the establishment of rational partitioning of electrons from the photosynthetic electron transport chain to hydrogen production by creation and implementation of in vivo PSI-Im7-hydrogenase fusions.

### Conclusion

In this study, we established and characterized Im7-E7 based bipartite (PSI-Fd) and tripartite (PSI-Fd-FNR) fusion systems for light-driven NADPH regeneration. In the bipartite PSI-Fd system, a 10 AA linker between E7 and Fd provided the highest electron transfer rate, most likely because this linker length allowed the required freedom and orientation between PSI-Fd and FNR. In the case of the tripartite PSI-Fd-FNR system, a 5 AA linker between Fd and FNR achieved the best electron transfer rate, which was somewhat lower than that of FPS, probably due to steric hindrance created by the two-sided tethering of Fd. Nevertheless, we believe that PSI fusions (either with or without Fd) based on the E7-Im7 system will have several advantages: i) it enables fusion of single or multiple oxidoreductases, ii) it can be implemented both in vitro and in vivo, iii) in vivo, it would enable channeling of electrons to specific oxidoreductase of interest by exploiting electrons from the photosynthetic electron transport chain.

## Material and Methods

### Strain construction

A *Synechocystis* PSI-PsaE-Im7-Histag mutant was generated using the plasmid pRSET6a_PsaE_Im7_Histag_Kan^R^ (Figure S10) via natural transformation. The PsaE gene together with flanking upstream and downstream regions (500 base pairs each) were amplified from the isolated genomic DNA of *Synechocystis* (primers 1&x2, Table S1) and inserted into an empty pRSET6a vector via an KpnI restriction site. Later a kanamycin resistance cassette (Kan^R^) was amplified from an external plasmid by PCR (primers 3&4, Table S1) and inserted in the downstream region also via restriction sites XmnI and Eco47III. The C-terminus of the *psaE* gene was extended by molecular cloning with Im7 gene, which was amplified from an pRSET6a_Im7_sfGFP plasmid by PCR (primers 5&6, Table S1) and inserted via the restriction site of SfoI. The transformation was performed as described (SI). Transformed *Synechocystis* mutant was screened, selected by growing on agar plates with 100 μg mL^-1^ kanamycin on agar plates. The homogeneity of the mutant strain was checked by PCR (primers 7&8, Table S1).

### Cultivation of *Synechocystis* and membrane preparation

*Synechocystis* cells were grown in a 25-L foil fermenter with BG11 medium at 30°C and bubbled with 5% (v/v) CO_2_-enriched air with final volume of 20L. The light intensity varied between 50-200 μE (white light) depending on the cell density. Cells were harvested at OD_680_ = 2 after three to five days of cultivation. The harvested, culture was concentrated to a volume of 2 L with a DC10 LA hollow fiber system (Amicon) and the cells were pelleted by centrifugation (25 °C, 20 min at 3,500 g). The pellet was resuspended in 80 mL of storage buffer 1 (20 mM MES pH 6.5, 10 mM MgCl_2_, 10 mM CaCl_2_, 500 mM mannitol and 20% (w/v) glycerol), and cells were snap-frozen in liquid nitrogen and eventually stored at −80 °C until further use. For membrane preparation, the cells were thawed, diluted with buffer A (20 mM MES pH 6.5, 10 mM MgCl_2_, 10 mM CaCl_2_), and pelleted (4 °C, 10 min at 21,000 g). Unless otherwise mentioned, all further steps were performed in dark or under green light illumination. The cell pellet was resuspended in 5 mL of buffer A supplemented with 0.4% (w/v) lysozyme per gram of fresh cell weight and incubated for 90 min at 37 °C. After centrifugation (4 °C, 15 min at 21,000 g), the pellet was resuspended in buffer A supplemented with 0.1 mg/mL of DNase (AppliChem) and 0.1 mM protease inhibitor (AEBSF Hydrochloride Biochemica, AppliChem) and subjected to a French press for cell disruption (1 min at 14,000 psi, four cycles). The crude thylakoid membrane pellet was collected by ultracentrifugation (4 °C, 30 min at 140,000 g) and resuspended in buffer A. Until further use, thylakoids were flash-frozen with liquid nitrogen and stored at −80 °C.

### Purification of PSI-Im7

Immobilized metal affinity chromatography was performed to isolate PSI-Im7. For this, thylakoid membranes were thawed, resuspended in extraction buffer (20 mM MES pH 6.5, 10 mM CaCl_2_, 10 mM MgCl_2_, 1.2% (w/v) DDM, 0.5% (w/v) sodium cholate) with a final Chl concentration of 1 mg/mL, and incubated for 30 min at 20 °C. ^[49,50]^ After ultracentrifugation (4 °C, 1 h at 140,000 g), NaCl and imidazole were added to the supernatant with final concentrations of 300 mM and 20 mM, respectively. The supernatant was filtered through a 0.45 μm syringe filter (Sarstedt AG) and applied to a 5 mL Ni Sepharose His trap crude FF column (GE-Healthcare), which was equilibrated with buffer B (20 mM MES pH 6.5, 10 mM CaCl_2_, 10 mM MgCl_2_, 500 mM mannitol, 300 mM NaCl, 0.03% (w/v) DDM and 20 mM imidazole). After washing the column with 50 mL of buffer B, bound PSI-Im7 was eluted by buffer C (20 mM MES pH 6.5, 10 mM CaCl_2_, 10 mM MgCl_2_, 500 mM mannitol, 300 mM NaCl, 0.03% (w/v) DDM and 500 mM imidazole). To concentrate the eluted fractions and to remove excess of NaCl and imidazole, a buffer exchange was performed using spin concentrators (Amicon, Ultra-15, 100 kD molecular weight cut off) with buffer D (20 mM MES pH 6.5, 10 mM CaCl_2_, 10 mM MgCl_2_, 500 mM mannitol, 0.03% (w/v) DDM). The collected PSI-Im7 complexes were flash-frozen with liquid nitrogen and stored at −80 °C until further use.

### SDS-PAGE-based characterization of PSI-Im7

PSI-Im7 subunit composition was analyzed by Tricine SDS-PAGE.^[51]^ Denatured subunits were separated in a separating gel containing 40% (w/v) polyacrylamide (acrylamide-bisacrylamide, 37.5:1) and 9.16 M urea, 4% (w/v) glycerol. Separating gel was overlaid with stacking gel of 8.1% polyacrylamide. PSI samples (equivalent to 2.5 μg Chl) were incubated in sample buffer for 30 min at 60 °C and loaded onto each lane. For all gel electrophoresis experiments, a PageRuler^™^ Prestained Protein Ladder (Thermo-Fischer-Scientific, product number 26616) was used. Electrophoresis was performed in Mini-PROTEAN System (BioRad) at 30 mA for 90 min with cathode buffer (0.1 M Tricine, 0.1% (w/v) SDS) in the gel-plate cassette and anode buffer (0.1 M Tris pH 8.9) in the electrophoresis chamber. The gels were stained by Coomassie staining solution (0.2% (w/v) Coomassie R250, 0.05% (w/v) Coomassie G250, 2% (v/v) ethanol, 5% (v/v) methanol, 10% (v/v) acetic acid After 30 min of incubation in staining solution, gels were de-stained using de-staining solution (7.5% (v/v) acetic acid and 5% (v/v) isopropanol).

### Characterization of the oligomerization state of PSI-Im7 by BN-PAGE

The oligomerization state of the isolated PSI-Im7 complex was analyzed by BN-PAGE. ^[52]^ Thereby, protein complexes were separated in a separating gel with a linear gradient of 3.2 to 16% polyacrylamide (acrylamide-bis-acrylamide, 32:1). PSI samples (equivalent to 3.5 μg Chl) were mixed with BN-PAGE loading buffer (50 mM Tricine, 15 mM Bis-Tris pH 7.0 with 50 % glycine) and loaded into each lane in the stacking gel. Gel electrophoresis was performed with a Mini-PROTEAN system (BioRad). Initially, electrophoresis was performed at 30 mA for 90 min with blue cathode buffer (15 mM Bis-Tris pH 7.0, 50 mM Tricine, and 0.02% (w/v) Coomassie Brillant blue 250). After 30 min, the gel chamber was exchanged with clear cathode buffer (15 mM Bis-Tris pH 7.0, 50 mM Tricine) and electrophoresis was continued at 150 V for additional 90 min.

### Determination of concentration of PSI-Im7 and PS1-Im7 fusion chimeras (PSI complexes)

The concentration of PSI complexes were determined based on chlrophyll amount. For extraction of chlorophyll, 1-2 μL of PSI complexes were dilluted in 1 mL methanol and the precipitated proteins of PSI complexes were pelleted by centrifugation (5 min, 21000 g). The chlorophyll amount was estimated by measuring absorbance of the supernatant at 652, 665 and 750 nm as previously described.^[53]^ Afterwards, the chlorophyll amount was converted into molar concentration by considering that each PSI monomer consists of 95 chlorophyll molecules.^[32]^

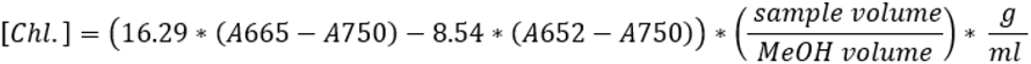

### ATR-FTIR difference spectroscopy

The surface of the germanium-IRE was functionalized as previously described.^[26]^ Briefly, the surface was cleaned, oxidized with a mixture of oxalic acid and hydrogen peroxide and silanized using the triethoxy linker-Silane molecule (2,5-dioxopyrrolidin-1-yl 4-oxo-4-((3- (triethoxysilyl) propyl) amino) butanoate). After silanization the crystal surface was functionalized with E7 (10 μg/ml in 50 mM HEPES buffer pH 7.5) and the remaining surface was blocked with casein. Subsequently, PS1-Im7 (10 μg/ml in buffer D) was immobilized on the surface. The binding of PSI-Im7 was followed spectrally using its amide II absorption kinetics (Figure S3). The obtained absorbance of 10 mAU corresponds to a surface density of about 1 pmol/cm^2^. For the measurement of the light induced difference spectrum of PSI-Im7, the sample compartment was darkened and a 250 W halogen lamp with fibre optic light conductor and long pass filter (> 645 nm) was installed in front of the cuvette. Subsequently, the immobilized PSI-Im7 was measured for 10 minutes during continuous illumination, followed by equilibration without light for 2 minutes and a measurement in darkness for 10 minutes. This measurement procedure was repeated ten times. For transmission measurement of PSI-Im7, 5 μg of protein in buffer D was added to an infrared cuvette with calcium fluoride windows (pathlength: 7μm). The cuvette was then placed in the optical path of the spectrometer. The measurement procedure with its illumination and dark cycles was then performed analogously to the ATR experiment. In Figure S4, the bands facing upwards are bands from the light adapted state and the bands facing downwards are from the dark state. Bands that do not change upon irradiation are cancelled out in this difference spectrum.

### Creation of chimeric E7-Fd and E7-Fd-FNR fusion proteins

A synthetic genetic construct of E7-Fd-FNR was designed *in silico* by a sequential assembly of three gene segments of E7, Fd, and FNR with additional sequences coding for AA linkers between the segments. Unique restriction sites were added at the beginning and end of each segment of the genetic construct (Figure S11), enabling easy replacement of individual segments. The genetic construct E7-Fd was created by removing the FNR segment of E7-Fd-FNR via restriction digestion with Bglll, Eco1ü9I. Additionally, a stop codon-containing oligonucleotide (nucleotide sequence 9&10, Table S1) was re-introduced at the end of the Fd segment by sticky-end ligation.

E7-Fd-FNR fusion chimeras were created by keeping linker lengths corresponding to 10 AA as non-variable between E7 and Fd segments, and linker length between Fd and FNR were varied by restriction digestion (BglII, XhoI) followed by sticky-end ligation with oligonucleotides (nucleotide sequences 11 to 14, Table S1) of respective linker length (5, 10, 15 AA). E7-Fd fusion chimeras were also created with variable linker lengths (5, 10, 15 AA) by restriction digestion (Hindlll, Ncol) followed by sticky-end ligation with the corresponding oligonucleotides (nucleotide sequences 15 to 18, Table S1). All genetic constructs were cloned and expressed in a pASK-IBA7 expression vector.

### Expression of E7-Fd and E7-Fd-FNR variants

For a pre-culture, 2 to 3 colonies of *E. coli BL21-ΔiscR* transformed with pASK-IBA7-E7-Fd or pASK-IBA7-E7-Fd-FNR were picked under sterile conditions and inoculated in 50 mL LB media supplemented with ampicillin (120 μg/mL) and grown overnight. 1 to 5% (v/v) of the starter culture was added to the main culture (LB media supplemented with 2 mM ferric ammonium citrate and 120 μg/mL of ampicillin) and incubated further at 37 °C under constant agitation at 130 rpm. As soon as the cultures reached an O.D_600_ of 0.6, protein expression was induced by the addition of 0.2 μg/mL of anhydrotetracycline and cultures were further incubated at 37 °C for 6 h. Then, cells were pelleted by centrifugation (4 °C, 20 min, 6842 g) and resuspended in TE buffer (10 mM Tris/HCl pH 8, 0.1 mM EDTA) to wash out the remnant growth medium.

### Isolation and purification of E7-Fd and E7-Fd-FNR variants

The harvested *E.coli* cells were suspended in equilibration buffer (100 mM Tris-HCl pH 8.0, 500 mM NaCl, 20 mM imidazole (3 mL/g of cell pellet) and treated with lysozyme (10 mg/ L^- 1^ cell culture), 0.1 % Triton X-100, 0.1 mM protease inhibitor (AEBSF Hydrochloride Biochemica, AppliChem) for 30 min. Cells were then subjected to sonication (on ice, 5×30 s with 45-60 s break between each pulse, 70 % duty cycle, output 4-5). After sonication, lysed cells were centrifuged (4 °C, 30 min, 76800 g) to remove insoluble particles and cell debris. The protein-containing supernatant was collected and filtered through a 0.45 μm size cut-off filter and applied onto a 5 mL His Trap^™^ crude FF column (GE Healthcare), running in equilibration buffer. The purification process was monitored and controlled by the ÄKTA FPLC-system. After washing with 30 ml of equilibration buffer, the His-tagged chimeric proteins were eluted by application of elution buffer (100 mM Tris-HCl pH 8, 500 mM NaCl, 500 mM imidazole). To concentrate the eluted fractions and to remove imidazole, a buffer exchange was performed using spin concentrators (Amicon, Ultra-15, 30 kD molecular weight cut off) with storage buffer 2 (100 mM Tris-HCl pH 8.0, 300 mM NaCl). Purified fusion proteins EF and EFF were also stored in storage buffer 2 at −80°C. The concentration of EF and EFF were estimated based on the extinction coefficients ε_422_ = 9.68 mM^-1^ cm^-1[54]^ and ε_459_ = 10.00 mM^-1^ cm^-1 [55]^, respectively. The expression and isolation protocols of other proteins used in this study, such as Fd, FNR, and *Cytc_6_* are mentioned in the supporting information (SI).

### UV-Vis spectra of E7-Fd and E7-Fd-FNR variants

The UV-Vis spectra of E7-Fd and E7-Fd-FNR fusion proteins were measured using a UV-2450 spectrophotometer (Shimadzu) and NanoDrop^™^ 2000 spectrophotometer (Thermofisher ^™^), respectively. An equivalent of 50 μM of chimeric E7-Fd fusion protein was used for spectral determination with a final volume of 1 mL in storage buffer 2. In the case of E7-Fd-FNR, to observe distinct peaks, a protein concentration of 0.4 mM was used with a final volume of 2 μL in storage buffer 2. Absorption spectra were recorded in a range between 200 and 700 nm and normalized to 280 nm. The extinction coefficient of the EF10 and E7 domain at 280 nm was calculated in the Expasy-ProtParam online tool based on protein sequence provided (SI).

### Light-induced NADP^+^ reduction

The reaction buffer used for the photoreduction of NADP^+^ in both bipartite and tripartite system was similar and adapted from previous literature.^[12]^ The reaction buffer was the following: 50 mM Tricine pH 8, 30 mM NaCl, 4 mM NaAsc, 100 μM DCPIP, 2.5 μM Cytc_*6*_, 5 mM MgCl_2_ (added only in the case of FPS1 or FPS2) and 1 mM NADP^+^. For the bipartite system 0.25 μM PSI-EF (5, 10, 15) chimeras and 0.75 μM free FNR were added to the reaction buffer. In case of FPS1 0.25 μM PSI, 0.25 μM Fd and 0.75 μM FNR were added to the reaction buffer. For the tripartite system 0.25 μM PSI-EFF (5, 10, 15) chimeras were added to the reaction buffer. In case of FPS2: PSI, Fd, and FNR were added in an equimolar ratio of 0.25 μM to the reaction buffer. The measurements were performed in a total volume of 1.5 mL at 25 °C. NADPH formation rates were analyzed by fluorescence spectroscopy (Dual-PAM-100, Walz Germany). The fluorescence was converted into NADPH quantity (μmol) based on the NADPH-fluorescence correlation curve (Figure S12). The total reaction time amounted to 8 min, whereby 1 min of pre-incubation in the dark was followed by a 5 min illumination with an actinic red light (100 μE) and a subsequent dark incubation for 2 min. The initial rate of NADP^+^ reduction was analyzed based on the slope obtained during the first min of illumination.

### Statistics and Software

All the experiments were performed in replicates to ensure the consistency and reproducibility of the data. Unless otherwise mentioned, all the data and graphs were processed and prepared with the software Origin 2019b.

## Abbreviations

Fd: ferredoxin
FNR: ferredoxin NADP^+^ reductase
E7: colicin E7 DNase
Im7: Immunity protein 7
NADP(H): (Dihydro) Nicotinamide Adenine Dinucleotide Phosphate
E7-Fd (EF): E7 DNase and ferredoxin fusion protein
E7-Fd-FNR (EFF): E7-DNase, ferredoxin, and ferredoxin NADP^+^ reductase fusion protein
PSI: Photosystem I
Cytc_6_: Cytochrome c_6_

## Acknowledgements

We acknowledge Dr. Kai Cormann for engineering the E7-Im7 system to utilize it as protein immobilization technique. We acknowledge Fabian Bisping for the creation of the PSI-PsaE-Im7-Histag mutant in *Synechocystis* sp. PCC. 6803 and we also acknowledge Dr. Nina Dyczmons-Nowaczyk for her help in designing the in silico E7-Fd-FNR construct. We thank Lucia Svoboda for her help in the optimization and characterization of PSI fusion chimeras and Jonas Simon for his help in the FTIR measurements. We acknowledge the financial support received from the European Union Horizon 2020 research and innovation program (grant agreement no.764920) as well as the Deutsche Forschungsgemeinschaft (DFG)-funded Research Training Group 2341 “Microbial Substrate Conversion (MiCon)”.

