## Supplementary Information for "A clickable photosystem I, ferredoxin, and ferredoxin NADP^+^ reductase fusion system for light-driven NADPH regeneration"

### Supplementary Text

#### Light-induced oxygen reduction with PSI-Im7 and PSI-Wt

The light-induced oxygen reduction of PSI was performed as described previously with slight adaptations [1]. The activity of PSI was measured based on light-induced Mehler reaction [2]. The oxygen reduction rate was monitored as a function of time as a functional parameter and measured by an optic oxygen sensor (FiBOX3, PreSens™). A PSI equivalent of 5 µg of chlorophyll (Chl) was diluted with PSI activity buffer (30 mM HEPES pH 7.5, 50 mM KCl, 3 mM $MgCl_2$, 330 mM mannitol, 0.3 % (w/v) β DDM, 5 mM sodium ascorbate, 0.5 mM methyl viologen, 0.8 mM DCPIP). Measurements were performed at 30 °C. After an initial 1 min dark incubation, PSI complexes were illuminated with red light (λ >665 nm, 4000 µmol $m^{-2}$ $s^{-1}$) for 4 min. The PSI-dependent oxygen reduction rate was calculated based on the slope of the light-induced reaction phase and specified as µmol $O_2$ mg $Chl^{-1}$ $mL^{-1}$ $h^{-1}$.

#### UV-vis spectrum of PSI-Im7 and PSI-Wt

Light absorption properties of isolated PSI-Wt and PSI-Im7 complexes were monitored by UV-Vis absorption spectra. Measurements were performed using a UV-2450 spectrophotometer (Shimadzu). PSI equivalent of 10 µg Chl was dissolved in 1 mL of buffer D (20 mM MES, pH 6.5, 10 mM $CaCl_2$, 10 mM $MgCl_2$, 500 mM mannitol, 0.03 % (w/v) β-DDM). Absorbance spectra were normalized at 680 nm.

#### Analyzing photoautotrophic growth of *Synechocystis* Wt and PSI-Im7

Photoautotrophic growth of *Synechocystis* sp. PCC 6803 (*Synechocystis* Wt) and the *Synechocystis* sp. PCC 6803 PasE-Im7 mutant (*Synechocystis* Im7) was monitored without antibiotics for 10 days under continuous light of 30 µmol $m^{-2}$ $s^{-1}$ at 30 °C and 150 rpm. The initial volume of cell cultures was 200 mL with an initial optical density at 750 nm ($OD_{750}$) of 0.1. The OD was measured periodically every second day for 10 days (n=4).

**Sucrose density gradient-based separation of excess E7-Fd and E7-Fd-FNR chimeras from PSI-Fd or PSI-Fd-FNR coupled complexes.**

For the preparation of the continuous sucrose gradient, 30 ml of a sucrose solution (50 mM HEPES pH 7.5, 10 mM $CaCl_2$, 10 mM $MgCl_2$, 0.03 % (w/v) β-DDM, 20 % (w/v) sucrose) was filled into SW-28 (Beckmann coulter) centrifuge tubes and incubated overnight at -20 °C. Later, the sucrose gradient was developed by thawing the solution for 2 h at room temperature. A 1:8 molar ratio of PSI-Im7 and fusion chimeras (EF, EFF) were mixed together and dark-incubated on ice for 30 min. Subsequently, a fusion mixture (~ 150 µg Chl) was applied on top of the gradient and subjected to ultracentrifugation (4°C, 18 h, 140000 g). The initial acceleration and deceleration speed of ultracentrifugation was adjusted to a low rotation speed (100 g). After ultracentrifugation, three distinct bands were observed: B1: EF or EFF fusion chimeras, B2: monomeric PSI coupled complex, and B3: Trimeric PSI coupled complex, as shown in Figure S7. B3 was collected carefully and washed with buffer D and flash-frozen with liquid $N_2$, and stored at -80 °C until further use.

**Genetic transformation of *Synechocystis* Wt**

50 mL of *Synechocystis* Wt were grown at 30 °C until $OD_{750}$: 0.5-1. The cells were harvested by centrifugation (15 min, 2900-3800 g) and resuspended in BG-11 medium up to a final cell concentration of $OD_{750}$: 2.0. 2.5 µg of plasmid DNA (pRSET6a_PsaE_Im7_Histag_Kan$^R$) were mixed with 0.4 mL of equivalent cell suspension by gentle shaking. Further, the cell culture was incubated under 50 µE standard light at 30 °C for 24 hours. Later, the cells were grown on a selective agar medium with a gradual increase in kanamycin concentration from initially 40 µg/mL to the final concentration of 100 µg/mL. This method was adapted from Williams, 1988[3].

**Fluorescence-based NADPH standard curve**

NADPH fluorescence was correlated to the concentration of NADPH. The calibration curve was created by adding the required NADPH in a stepwise manner. 1.4 mL of reaction buffer (50 mM Tricine pH 8, 30 mM NaCl, 5 mM $MgCl_2$) was added to a quartz cuvette and 5 µL of 284.03 µM NADPH stock solution was added in subsequent steps so that concentrations from 0 to 20 µM NADPH were achieved. The mixture was stirred briefly with a magnetic stirrer and allowed to settle for 1 min. Subsequently, the fluorescence of the corresponding NADPH concentration was measured for 30 s. The measurement was performed in triplicates for each concentration. The mean fluorescence value corresponding to each NADPH concentration was graphically represented, and the slope was calculated in Orgin19b. The calculated slope was 0.137 arbitrary units (a.u.) / µM NADPH.

##### Nucleotide sequences for linker variation

The nucleotide sequences for linker variation were created using oligonucleotides of respective lengths. Oligonucleotides were designed in Clone Manager 9 and were ordered from Sigma-Aldrich Ltd. Two complementary sequences of individual linker lengths (Table S1) were annealed by exposing sticky ends. For annealing, two complementary oligonucleotides: 4.5 µM forward and 4.5 µM reverse oligonucleotides of respective linker lengths (5, 10, 15 amino acids), were mixed with 1x Taq buffer (Thermo Scientific Ltd.) and 180 mM NaCl. The total volume was set to 30 µL with $H_2O$. Subsequently, the oligonucleotide mixture was incubated at 100 °C for 5 min and allowed to cool down slowly to room temperature. Later, excess Taq buffer and NaCl was removed from the annealed oligonucleotides via alcohol precipitation. Finally, precipitated oligonucleotides were resolved in TE buffer (10 mM Tris-HCl, pH 8.0, 1 mM EDTA) and stored at –20 °C until further use.

##### Heterologous recombinant expression of Fd (PetF)

2 to 3 colonies of *E. coli* BL21 Δ*iscR* cells [4] transformed with pASK-IBA7-PetF-streptag (PetF sequence from *T. vestitus* BP1) were grown on an agar plate supplemented with 120 µg/mL ampicillin and were picked under sterile conditions to inoculate 50 mL LB media supplemented with antibiotics (120 µg/mL ampicillin) as preculture. 15 mL of preculture was added to the 1 L main culture (LB media + 2 mM ferric ammonium citrate) supplemented with antibiotics. Culture growth was monitored by absorbance at $OD_{600}$. After reaching an $OD_{600}$ of 0.4-0.6, the culture was induced for protein expression with 0.2 µg/mL of anhydrotetracycline. After incubation of the culture for 6 h at 37 °C, cells were pelleted by centrifugation (4 °C, 20 min, 6900 g) and washed with TE buffer (10 mM Tris-HCl pH 8.0, 1 mM EDTA) for the removal of growth medium residues.

**Isolation and purification of Fd (PetF)**

The harvested *E. coli* cells were suspended in streptag equilibration buffer (100 mM Tris-HCl pH 8.0, 3 mL/g of cell pellet) and treated with lysozyme (10 mg /$L^{-1}$ cell culture), 0.1 % (v/v) Triton X-100, 0.1 mg/mL of DNase (AppliChem), and 0.1 mM protease inhibitor (AEBSF Hydrochloride Biochemica, AppliChem) for 30 min. Further, the sample was subjected to sonication (on ice, 5×30 s with 45-60 s break between each pulse, 70 % duty cycle, and output between 4-5). After sonication, the sample was centrifuged (4 °C, 30 min, 21000 g) to remove insoluble particles and the collected supernatant was filtered through a 0.45 µm cut-off filter. Later, the supernatant was applied on to a 5 mL Strep-tag column (GE Healthcare) pre-equilibrated with streptag equilibration buffer for purification of Fd., The Strep-tag column bound Fd was eluted by streptag elution buffer (100 mM Tris-HCl pH 8.0, 4 mM desthiobiotin). The purification process was monitored and controlled by the ÄKTA FPLC™-system. To concentrate the eluted fractions and to remove desthiobiotin, a buffer exchange was performed using spin concentrators (Amicon, Ultra-15, 3 kDa molecular weight cut off) with storage buffer 2 (100 mM Tris-HCl pH 8.0, 300 mM NaCl) and purified Fd was stored in storage buffer 2 at -20 °C. Protein concentration of Fd was calculated via absorbance at 422 nm using the molar absorption coefficient $\varepsilon = 9.8\ mM^{-1}\ cm^{-1}$[5].

**Heterologous recombinant expression and purification of FNR (PetH)**

2 to 3 colonies of *E. coli* BL21 transformed cells with pASK-IBA37$^{+}$-PetH-Histag (PetH sequence from *T. vestitus* BP-1) picked under sterile conditions to inoculate 50 mL LB media supplemented with antibiotics (120 µg/mL ampicillin) as preculture. 15 mL of preculture was added to the 1 L main culture (LB media) supplemented with antibiotics. After reaching an $OD_{600}$ of 0.4-0.6, the culture was induced for protein expression with 0.2 µg/mL of anhydrotetracycline and further incubated the culture for 6 h at 37 °C. Later, the cells were pelleted by centrifugation (4 °C, 20 min, 6900 g) and washed with TE buffer (10 mM Tris-HCl pH 8.0, 1 mM EDTA) for the removal of growth medium residues. For purification, the FNR overexpressed *E. coli* cells were suspended in equilibration buffer (100 mM Tris-HCl pH 8.0, 500 mM NaCl, 20 mM Imidazole (3 mL/g of cell pellet). The suspended cells were treated with lysozyme (10 mg/ $L^{-1}$ cell culture), 0.1 % Triton X-100, 0.1 mM protease inhibitor (AEBSF Hydrochloride Biochemica, AppliChem) for 30 min. Cells were then

subjected to sonication on ice (5×30 s with 45-60 s break between each pulse, 70 % duty cycle, and output between 4-5). After sonication, the sample was centrifuged (4 °C, 30 min, 21000 g) to remove insoluble particles and the collected supernatant was filtered through a 0.45 µm cut-off filter. Later, the protein-containing supernatant was collected and filtered through a 0.45 µm size cut-off filter and applied onto 5 mL His Trap™ crude FF column (GE Healthcare) for the purification. The His-tagged chimeric proteins were eluted by application of elution buffer (100 mM Tris-HCl pH 8, 500 mM NaCl, 500 mM imidazole) in a gradient manner onto the column. The purification process was monitored and controlled by the ÄKTA FPLC™-system. To concentrate the eluted fractions and to remove imidazole, a buffer exchange was performed using spin concentrators (Amicon, Ultra-15, 30 kDa molecular weight cut off) with storage buffer 2 and purified FNR was stored in storage buffer 2 at -20 °C. Protein concentration of FNR was calculated via absorbance at 459 nm using the molar absorption coefficient ε =10.00 $mM^{-1}$ $cm^{-1}$[6].

**Heterologous expression and purification of Cytc$_6$ (PetJ)**

pASK-IBA4-PetJ (PetJ sequence from *T. vestitus* BP-1)-Histag-transformed *E. coli* BL21-PEC86 cells were grown on an agar plate supplemented with 120 µg/mL ampicillin and 30 µg/mL chloramphenicol. For the preparation of a preculture, 2-3 transformed colonies were picked under sterile conditions and inoculated in 50 mL LB media supplemented with 120 µg/mL ampicillin, 30 µg/mL chloramphenicol. A 15 mL of preculture was added to 1 L of the main culture (TB media). After reaching an $OD_{600}$ of 0.8, the culture was induced for protein expression with 0.2 µg/mL of anhydrotetracycline and transferred into screw neck laboratory glass bottles to achieve microaerobic conditions for high yield expression of Cytc$_6$. The culture was incubated under microaerobic condition for 12 h at 25 °C with continuous agitation at 100 rpm. After incubation, the cells were harvested and stored as mentioned above in Fd and FNR recombinant expression procedure. Unless otherwise specified, the purification of Cytc$_6$ was performed as described for FNR. To concentrate the eluted fractions and to remove imidazole, a buffer exchange was performed using spin concentrators (Amicon, Ultra-15, 3 kDa molecular weight cut off) with storage buffer 2 and purified Cytc$_6$ was stored in storage buffer 2 at -20 °C. . The concentration of the Cytc$_6$-Histag was estimated via absorption at 552 nm using the molar absorption coefficient ε=24.4 $mM^{-1}$ $cm^{-1}$.

### Supplementary Figures

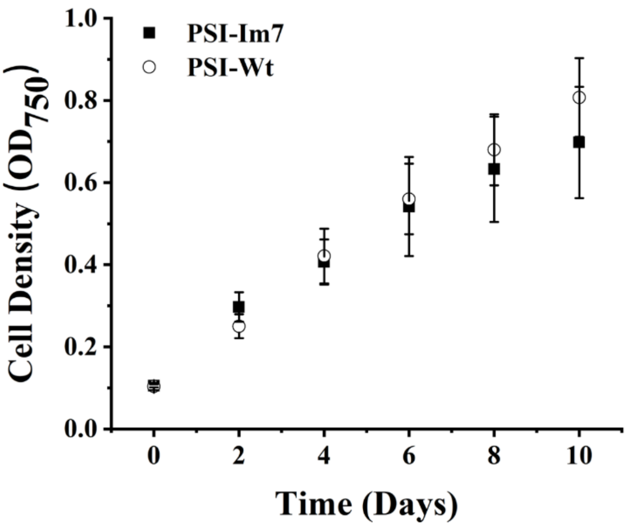

**Supplementary Figure 1: Photoautotrophic growth of *Synechocystis* Wt (open circle) and *Synechocystis* PSI-Im7 mutant (dark square).** The photoautotrophic growth was monitored by measuring optical density at 750 nm. Data represent the mean values and errors bars indicate the standard deviation of independent biological replicates (n=4).

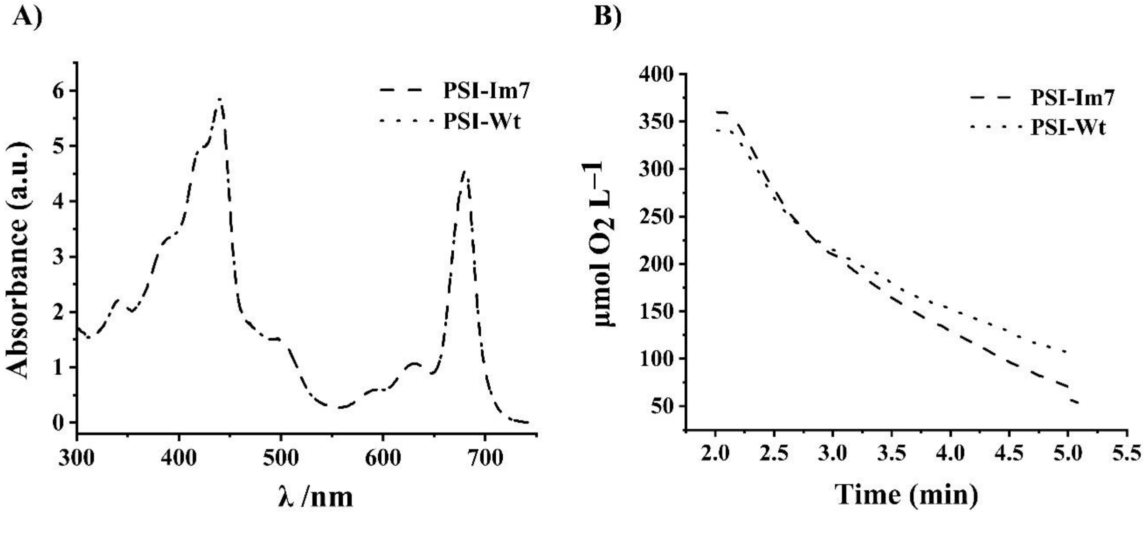

**Supplementary Figure 2**. **Oxygen reduction rate and UV-Vis spectral characterization of PSI-Im7 (Dashed line) & PSI-Wt (Dotted line).** A) UV-Vis absorption spectra of isolated PSI-Im7 and PSI-Wt. The spectral range was set to 300 to 750 nm and the data was normalized to the absorbance at 680 nm. Measurements were carried out in triplicates and the data represent the mean value. B) The oxygen reduction rate is normalized to the chl *a* concentration. The total measurement time was 5 min with a dark (1 min)- light (4 min) cycle. Measurements were carried out in triplicates and the mean value is shown for each data point.

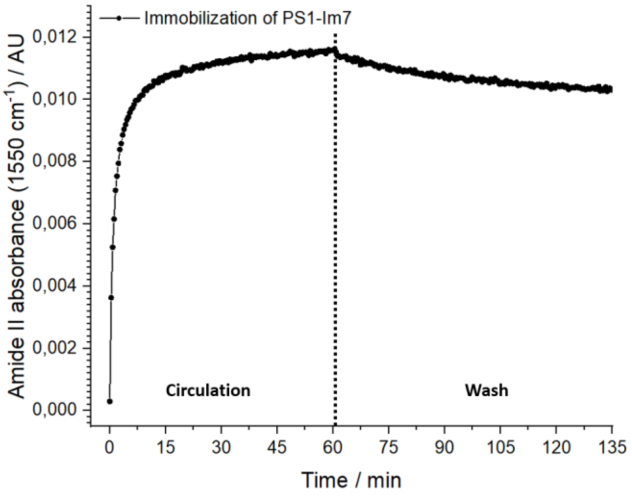

**Supplementary Figure 3**: Kinetic profile of the PSI-Im7 immobilization on the E7-modified germanium surface. Shown is the time course of amide II absorption at 1550 cm$^{-1}$. The dashed line highlights the switch from the circulation step with PS1-Im7 to the washing step with buffer D.

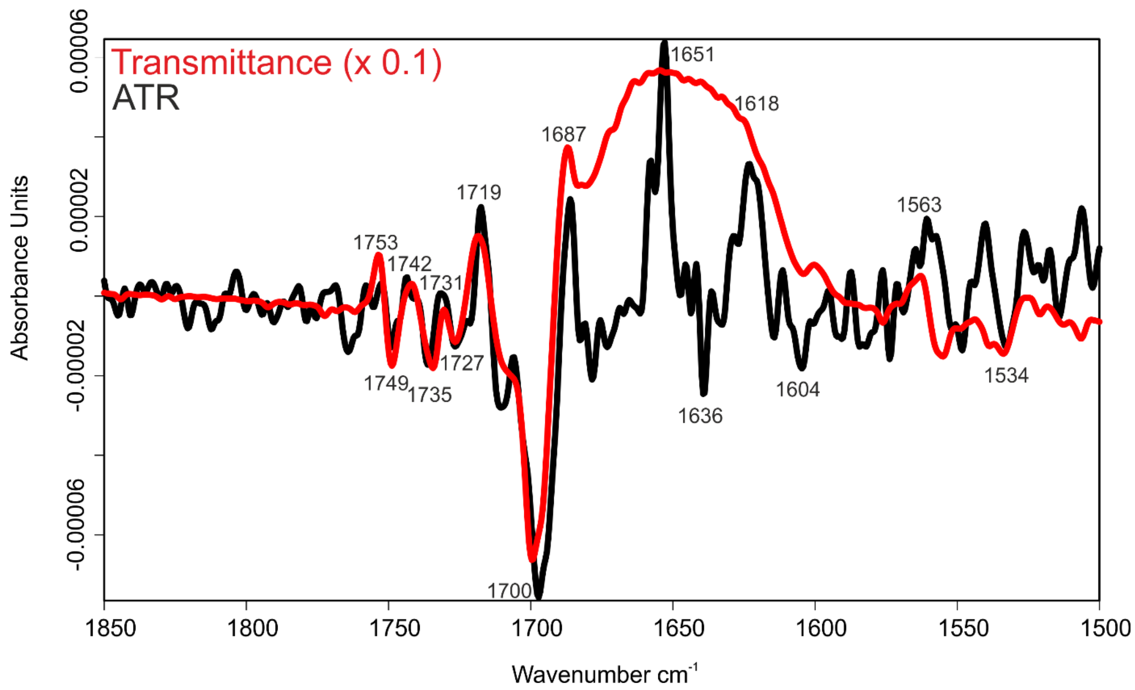

**Supplementary Figure 4**: IR-difference spectra of the light-induced response of PSI-Im7. The black spectrum results from the ATR measurement of immobilized PS1-Im7 on the E7-modified germanium crystal. The red spectrum was obtained by a transmission measurement. The latter one is obscured in the range between 1680 cm$^{-1}$ and 1600 cm$^{-1}$ due to the total absorption of water.

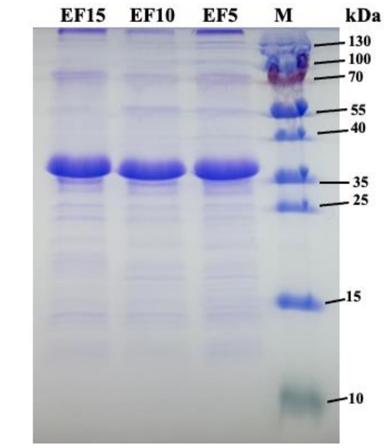

**Supplementary Figure 5. SDS-PAGE-based analysis of E7-Fd (EF) chimeras purified by His-tag affinity chromatography.** From left to right**:** Lanes 1 to 3: EF chimeras, with amino acid linker lengths of 5, 10, and 15 amino acids, respectively. Lane 4: Protein marker (PageRuler, Thermoscientific$^{TM}$). An equivalent of 10 µg protein was loaded onto each lane.

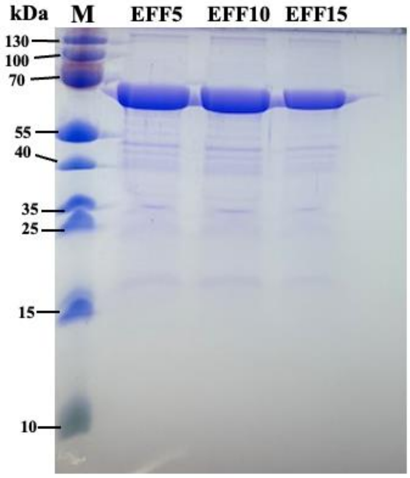

**Supplementary Figure 6. SDS-PAGE analysis of E7-Fd-FNR (EFF) chimeras purified by His-tag affinity chromatography.** From left to right: Lane 1: Protein marker (PageRuler, Thermoscientific$^{TM}$). Lanes 2 to 4: EFF chimeras, with linker length variations of 5, 10, 15 amino acids between Fd and FNR, respectively. An equivalent of 10 µg protein was loaded onto each lane.

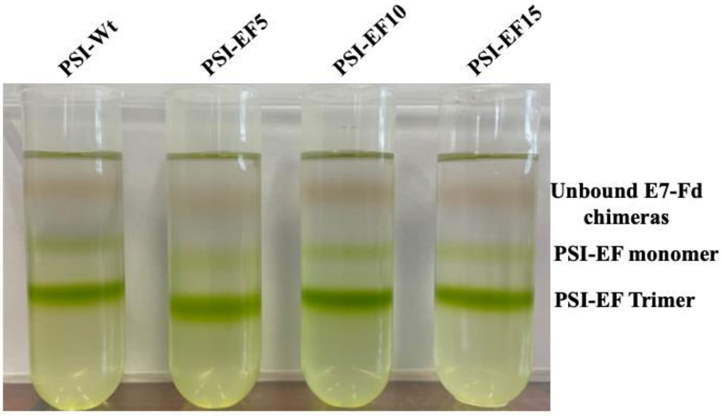

**Supplementary Figure 7**. **Continuous sucrose density gradient centrifugation PSI-EF chimeras**. Separation of unbound EF chimeric from PSI-Im7-coupled E7-Fd chimeras on a linear 0-20 % (w/v) sucrose density gradient. A mixture equivalent to 150 µg Chl and corresponding molar concentrations of EF were loaded onto each centrifugation tube.

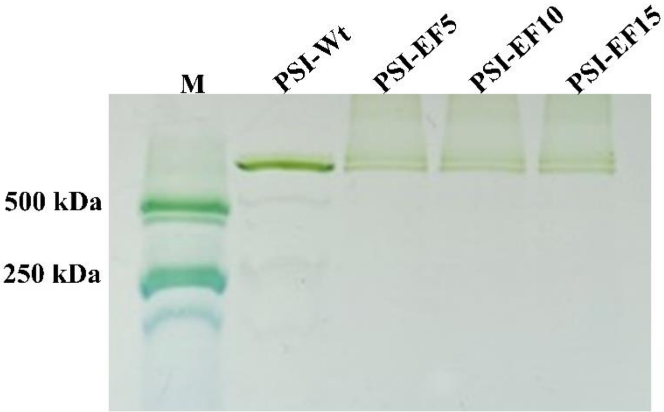

**Supplementary Figure 8**. **BN-PAGE-based analysis of PSI-Fd (PSI-EF) chimeras. From Left to right:** Lane 1: Photosystem II (PSII) dimer (500 kDa) and monomer (250 kDa) from *T. vestitus* BP-1, Lane 2: PSI-Wt from *Synechocystis*(trimeric) after incubation with EFF5, Lane 3 to 5: PSI-Im7 (trimeric) after coupling with EF chimeras (5 to 15), respectively. Equivalents of 1 µg Chl PSI fusion complexes were loaded onto each lane.

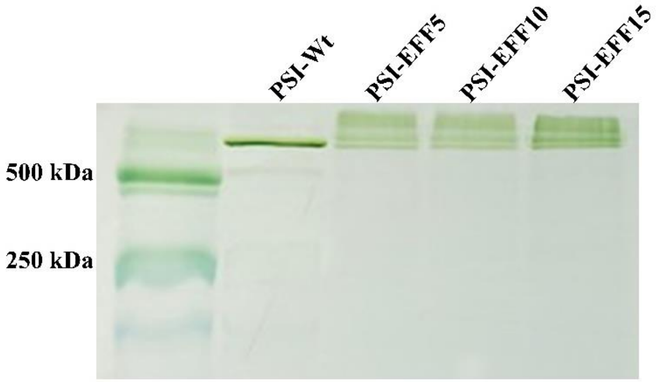

**Supplementary Figure 9. BN-PAGE of PSI-Fd-FNR (PSI-EFF) chimeras.** Lane 1: PSII dimer (500 kDa) and monomer (250 kDa) from *T. vestitus* BP-1, Lane 2: PSI-Wt from *Synechocystis* (trimeric) after coupling with EFF5, Lane 3 to 5: PSI-Im7 (trimeric) after coupling with EFF chimeras (5 to 15, respectively). PSI equivalents of 1 µg Chl were loaded onto each lane.

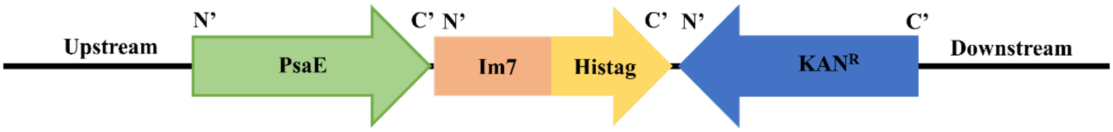

**Supplementary Figure 10. Schematic representation of the genetic construct of plasmid pRSET6a_PsaE_Im7-Histag_KAN$^R$ used for the creation of the *Synechocystis* PSI-Im7 mutant.**

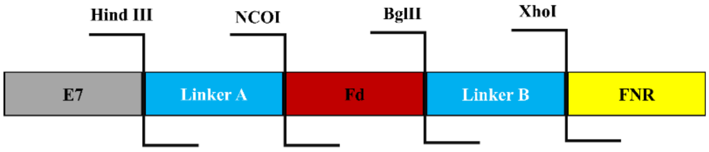

**Supplementary Figure 11. Schematic representation of synthetic genetic construct of E7-Fd-FNR fusion protein.** Each gene segment is separated with a nucleotide sequence corresponding to the amino acid linker. The end of each gene segment consists of a unique restriction site.

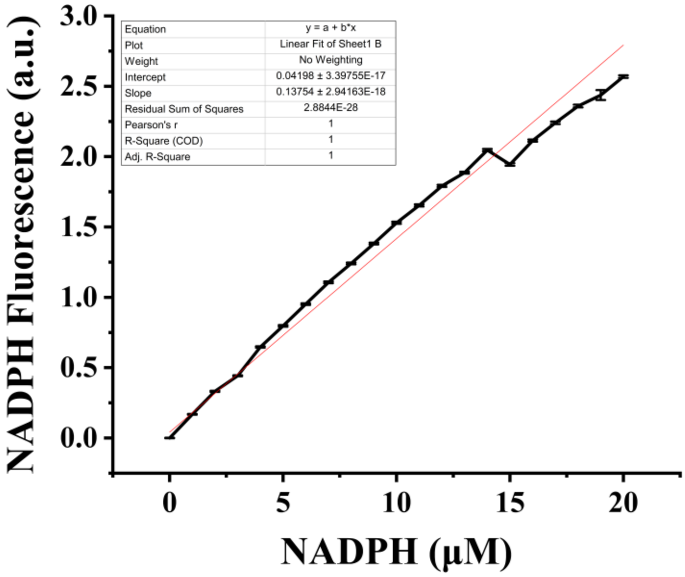

**Supplementary Figure 12. Fluorescence-based NADPH calibration curve.** The fluorescence of the corresponding NADPH concentration from 0 to 20 μM was measured. The data represents the mean values of fluorescence corresponding to each NADPH concentration and the standard deviation is represented with error bars (n=3).

```
DNase E7            1 ESKRNKPGKATGKGKPVNNKWLNNAGKDLGSPVPDRIANKLRDKEFKSFD     50
                      |||.||||||||.|.|||||||||||:||||||||||||.||.:||.|||
DNase_Opti_E7       1 ESKENKPGKATGDGDPVNNKWLNNAGEDLGSPVPDRIANDLRCEEFDSFD     50

DNase E7           51 DFRKKFWEEVSKDPELSKQFSRNNNDRMKVGKAPKTRTQDVSGKRTSFEL    100
                      ||||||||||||||||||||||||||||||||||||||||||||||||||
DNase_Opti_E7      51 DFRKKFWEEVSKDPELSKQFSRNNNDRMKVGKAPKTRTQDVSGKRTSFEL    100

DNase E7          101 HHEKPISQNGGVYDMDNISVVTPKRHIDIHRGK    133
                      |.|||||||||||||||||||||||||||||||
DNase_Opti_E7     101 HAEKPISQNGGVYDMDNISVVTPKRHIDIHRGK    133
```

**Supplementary Figure 13. DNA sequence alignment of E7-DNase and E7-DNase_Opti.** Comparative analysis of semi-conserved and conserved substitution of amino acid in E7-DNase_Opti. The alignment was made using EMBOSS matcher.

### Supplementary Table

| **Supplementary Table 1: Oligonucleotides and primer used to create plasmid constructs (pASK-IBA7-E7-Fd-(FNR), PRSET6a-PsaE_Im7_Histag_Kan$^R$).** | | | |
|---|---|---|---|
| **No** | **Name of Linker** | **DNA sequence of linker 5'→ 3'** | **Purpose** |
| 1 | PsaE-CT-forward | CAAGTTTTGGGAGTAGGGGAGC | Used for amplification of PsaE along with upstream and downstream sequence from isolated *Synechocystis* Wt genome |
| 2 | PsaE-CT-reverse | GGGCATTTTTAAAGGGCGGCG | Used for amplification of PsaE along with upstream and downstream sequence from isolated *Synechocystis* Wt genome |
| 3 | KanR-forward | CCGCTCATGAATTAATTCTTAGAAAAACTC ATCG | Used for amplification of Kanamycin resistance cassette from plasmid PRSF-Duet1 |
| 4 | KanR-reverse | GAGCGCTGCATGCCTATTTG | Used for amplification of Kanamycin resistance cassette from plasmid PRSF-Duet1 |
| 5 | Im7-forward | GGTGGGTCTGGTGAACTGAAAAATAGTATT AGTGATTACACAGAGGC | Used for amplification of Im7 gene from pRSET6a_Im7_sfGFP plasmid |
| 6 | Im7-reverse | TTAATGGTGATGGTGATGGTGATGGTGATG GTGAGCGCCTCCGCTGCCGCC | Used for amplification of Im7 gene from pRSET6a_Im7_sfGFP plasmid |
| 7 | PsaE-Im7-forward | GATATCGCCCGCTCATGAATTAATTCTTAG | Used for segregation check of PSI_Im7 mutant |
| 8 | PsaE-Im7-reverse | TTTTGCCGCCGCTTG | Used for segregation check of PSI_Im7 mutant |
| 9 | Fd-stop reverse | GATCTTAGCTCGAGTGGCGGTGG | For the creation of E7-Fd construct from E7-Fd-FNR, a stop codon was reintroduced at the end of Fd with this oligonucleotide |
| 10 | Fd-stop forward | GCCCCACCGCCACTCGAGCTAA | For the creation of E7-Fd construct from E7-Fd-FNR, a stop codon was reintroduced at the end of Fd with this oligonucleotide |
| 11 | B5 reverse | TCGAGACTACCACTACCGCCA | Linker exchange to 5 AA between Fd and FNR |
| 12 | B5 forward | GATCTGGCGGTAGTGGTAGTC | Linker exchange to 5 AA between Fd and FNR |
| 13 | B15 reverse | TCGAGACTACCGCCACTCCCGCCACTGCCT CCGCTACCTCCGCTCCCACCA | Linker exchange to 15 AA between Fd and FNR |
| 14 | B15 forward | GATCTGGTGGGAGCGGAGGTAGCGGAGGC AGTGGCGGGAGTGGCGGTAGTC | Linker exchange to 15 AA between Fd and FNR |
| 15 | A5 reverse | CATGGACTACCACTCCCGCCA | Linker exchange to 5 AA between E7 and Fd |
| 16 | A5 forward | AGCTTGGCGGGAGTGGTAGTC | Linker exchange to 5 AA between E7 and Fd |
| 17 | A15 reverse | CATGGACTACCGCCACTCCCGCCACTGCCT CCGCTACCTCCGCTCCCACCA | Linker exchange to 15 AA between E7 and Fd |
| 18 | A15 forward | AGCTTGGTGGGAGCGGAGGTAGCGGAGGC AGTGGCGGGAGTGGCGGTAGTC | Linker exchange to 15 AA between E7 and Fd |

### Protein sequences of E7-Fd, E7-Fd-FNR, PsaE-Im7-Histag

#### Protein sequences of EF

MHHHHHHASESKENKPGKATGDGDPVNNKWLNNAGEDLGSPVPDRIANDLRCEEFDSFDDFRKK
FWEEVSKDPELSKQFSRNNNDRMKVGKAPKTRTQDVSGKRTSFELHAEKPISQNGGVYDMDNISV
VTPKRHIDIHRGKKLGGSGGSGGSGPWATYKVTLVRPDGSETTIDVPEDEYILDVAEEQGLDLPFSC
RAGACSTCAGKLLEGEVDQSDQSFLDDDQIEKGFVLTCVAYPRSDCKILTNQEEELYRS

| | |
|---|---|
| Linker 5 aminoacids | GGSGG |
| Linker 10 aminoacids | GGSGGSGGSG |
| Linker 15 aminoacids | GGSGGSGGSGGGSGG |

#### Protein sequence of EFF

MHHHHHHASESKENKPGKATGDGDPVNNKWLNNAGEDLGSPVPDRIANDLRCEEFDSFDDFRK
KFWEEVSKDPELSKQFSRNNNDRMKVGKAPKTRTQDVSGKRTSFELHAEKPISQNGGVYDMDNI
SVVTPKRHIDIHRGKKLGGSGGSGGSGPWATYKVTLVRPDGSETTIDVPEDEYILDVAEEQGLDLP
FSCRAGACSTCAGKLLEGEVDQSDQSFLDDDQIEKGFVLTCVAYPRSDCKILTNQEEELYRSGGSG
GSGGSGLEYNATNSRSRMFRYEVVGLRQTAETEKTNYAIRNSGSQFFNVPYDRMNQFMQQITRW
GGKIVSIQPLNGTVAPLAATTEPAANNGAAPVKEKKVDIPVNIYRPNNPCIGKVISNEELVREGGE
GTVKHIIFDISGTELRYLEGQSIGIIPAGTDANGKPHKLRLYSIASTRHGDFQDDKTVSLCVRRLEY
KDKETGETIYGVCSSYLNQLQPGDEVKITGPVGKEMLLSDDPEATIIMLATGTGIAPFRAFLWRMF
KENNPDYQFKGLAWLFFGVAYTANILYKDELEAIQAQYPDHFRLTYAISREQKTPDGGKMYIQGR
IAEHADEIWQLLQKKNTHVYMCGLRGMEPGIDEAMTAAAAKNGADWQEFLKGTLKKEGRWHV
ETY

| | |
|---|---|
| Linker 5 aminoacids | GGSGG |
| Linker 10 aminoacids | GGSGGSGGSG |
| Linker 15 aminoacids | GGSGGSGGSG GGSGG |

#### Protein sequence of PSaE-Im7-Histag

MALNRGDKVRIKRTESYWYGDVGTVASVEKSGILYPVIVRFDRVNYNGFSGSASGVNTNNFAEN

ELELVQAAAKGGSGELKNSISDYTEAEFVQLLKEIEKENVAATDDVLDVLLEHFVKITEHPDGTDL

IYYPSDNRDDSPEGIVKEIKEWRAANGKPGFKQGGSGGAHHHHHHHHHH
